# Threat-dependent scaling of prelimbic dynamics to enhance fear representation

**DOI:** 10.1101/2023.10.10.561506

**Authors:** José Patricio Casanova, Clément Pouget, Nadja Treiber, Ishaant Agarwal, Mark Allen Brimble, Gisella Vetere

## Abstract

Promptly identifying threatening stimuli is crucial for survival. Freezing is a natural behavior displayed by rodents towards potential or actual threats. While it is known that the prelimbic cortex (PL) is involved in both risk evaluation and in fear and anxiety-like behavior expression, here we explored whether PL neuronal activity can dynamically represent different internal states of the same behavioral output. We found that freezing can always be decoded from PL activity at a population level. However, the sudden presentation of a fearful stimulus quickly reshaped PL to a new neuronal activity state, an effect not observed in other cortical or subcortical regions. This shift changed PL freezing representation and is necessary for fear memory expression. Our data reveal the unique role of PL in detecting threats and internally adjusting to distinguish between different freezing-related states.

**IN BRIEF:** Through a comparative analysis across brain regions and risk situations, we demonstrate the distinctive role of the Prelimbic Cortex (PL) in fear-related behavior representation: the PL promptly detects threat stimuli and adjusts its neuronal configuration accordingly, resulting in a change in freezing representation, which is necessary for memory-associated fear expression.

**HIGHLIGHTS:** - PL population activity represents fear-related behavior (i.e., freezing)

- Same behavioral outcome (freezing) can be represented as different PL neuronal states

- Other brain regions do not show changes in dynamics to represent freezing

- PL dynamics following threats are necessary for proper fear memory expression

## INTRODUCTION

Recognizing and responding to potentially dangerous stimuli requires monitoring of information from external cues and internal states. A network of brain regions, with the medial prefrontal cortex (mPFC) as a central hub, is believed to be engaged in this task. The mPFC is in a privileged position ^1^, to receive, integrate and compute sensory information, drive output circuits, and organize behavior ^2,3^. The correct recruitment of the Prelimbic (PL) subdivision of the mPFC is necessary for evaluating and modulating fear and anxiety-like behavior ^4–12^. Neuronal dynamics must maintain a balance between flexibility, to cope with an ever-changing environment, and stability, to retain relevant information ^13^. mPFC shows neuronal flexibility in response to threatening experiences and adjustment to engage in avoidance behavior ^14^. However, mice can employ various strategies in response to identical threats. In contextual fear conditioning, mice exhibit a passive defense strategy (i.e., freezing behavior), following inescapable foot shock delivery. Whether the PL modulates its state to represent freezing in response to a threat is unknown. Although freezing is interpreted as an indicator of fear, it can be displayed in situations where no immediate threat is present. In these instances, freezing may reflect the mice’s anxiety or stress, while evaluating the risk. Freeze responses can be elicited by various stimuli including novel environments or objects ^15^. While freezing is observable, it can correspond to different internal states in mice. Here we investigate whether the presentation of a threat alters the dynamics of the PL, resulting in distinct internal representations of freezing. We tested mice in 3 tasks that elicited freezing behavior while monitoring PL population activity with in vivo calcium imaging. In a different group of mice we also imaged from the Retrosplenial cortex (RSC) and a subcortical region, the Laterodorsal Thalamus (LDn) and compared with PL results, providing insights into the differential neural mechanisms underlying risk evaluation and subsequent fear representation.

## RESULTS

### PL single cells respond to freezing and threat and show memory signatures 1 day after

PL neuronal activity was imaged while mice were tested in 3 behavioral tasks eliciting freezing behavior (Fig 1A). Representative occupancy heatmaps in every task are shown in Fig 1B. In all tasks, we quantified time spent in freezing (Fig 1C, Fig S1A). In the Novel Object (NO) task, mice were first re-exposed to the same context of OF for 3 minutes and after 3 minutes, we introduced a novel object ^16^. We found an increase in freezing after the presentation of the novel object (Fig S1B, W_(10)_=49, P<0.01, for detailed statistics see Supplementary Table 1), and overall freezing increased compared to the OF (Fig 1C; F_(1.381, 14.73)_=35,07, P<0.0001; OF vs NO, P<0.0021, Tukey’s post-hoc test). During Contextual Fear Conditioning (CFC), mice received 2 foot shocks (FS). Freezing score in CFC was similar to the OF (Fig 1C, P=0.5523, Tukey’s post-hoc test), suggesting that mice were displaying behavioral responses associated with exploring equivalently novel contexts. However, when comparing the 2 minutes before versus after FS, we observed an increase in freezing (Fig S1C, W_(9)_=41, P<0.05). One day after CFC, mice were re-exposed to the conditioning context (CFTEST). Here, mice displayed the highest freezing, compared to every other task (Fig 1C, CFTEST vs OF, P<0.01; CFTEST vs NO, P<0.05; CFTEST vs CFC, P<0.0001, Tukey’s post-hoc test). These data show that the tasks elicited freezing responses of different magnitudes towards potential or actual threats, confirming it can be adopted in a variety of situations, including anxiety and risk evaluation ^15,17^.

**Figure 1.**
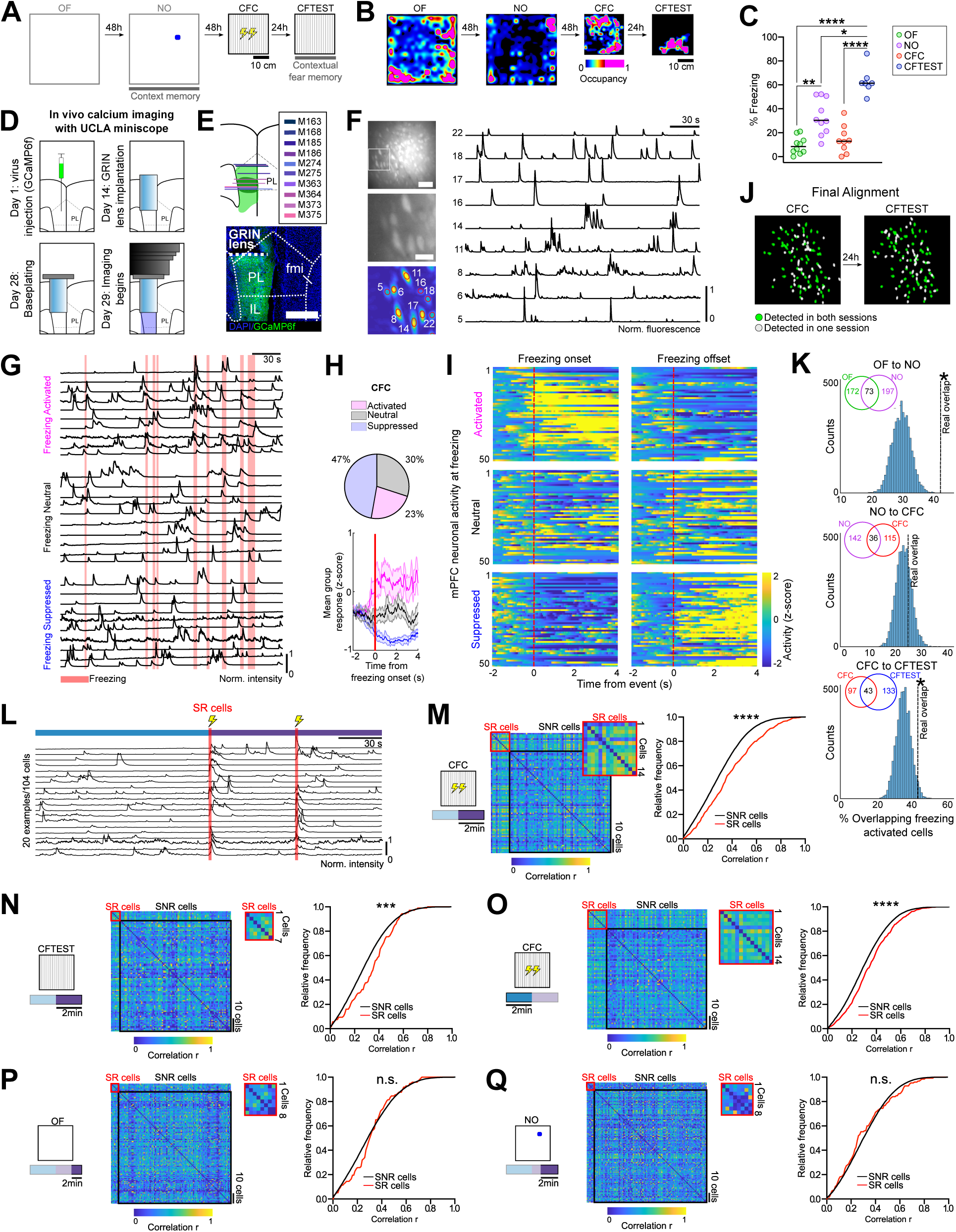
PL single neurons dynamically encode freezing and FS information across different experiences. **(A)** Time course of behavioral tests. **(B)** Representative occupancy heat maps. **(C)** Freezing scores. Black bars represent the median. **(D)** Timeline for in vivo calcium imaging. **(E)** Top, schematics of GRIN lens placement in the PL. Green areas represent minimum (dark) and maximum (light) GCaMP6f virus spread. Bottom, GRIN lens aimed at the PL. Scale bar 500 µm. **(F)** Maximum projection from raw miniscope field of view (top left, scale bar 50 µm) and magnification (middle left, scale bar 20 µm). Bottom left, identified neurons ROIs. fmi, forceps minor of the corpus callosum; IL, Infralimbic cortex. Right, representative calcium traces. **(G)** Calcium traces from freezing activated (top), freezing neutral (middle), and freezing suppressed cells (bottom) during CFC. **(H)** Top, PL cells classification towards freezing during CFC (N=866 neurons). Bottom, mean group response to freezing onset (red line). Data represent mean ± SEM. **(I)** Heatmaps showing normalized activity of 50 cells aligned to freezing onset and offset. **(J)** Example of cell registration of cells across tests. **(K)** Percentage of overlapping freezing-activated cells vs a null distribution. Inset, number of freezing-activated cells detected at pairs of consecutive tests. **(L)** Representative SR cells (total N=104 cells). **(M-Q)** Normalized fluorescence correlation matrices and relative frequency plots of the correlation between SR (red) or SNR (black) cells during CFC, following FS **(M)**, during CFTEST **(N)**, during CFC before the first FS **(O),** during the last two minutes of OF **(P)** and NO **(Q)**. Each matrix represents data from the same representative mouse. Insets, zoomed SR correlation matrices. *P<0.05, **P<0.01, ***P<0.001, ****P<0.0001.

We monitored PL activity underlying freezing from individual neurons with a miniaturized head-mounted fluorescence microscope ^18,19^. We injected an adeno-associated virus (AAV) packaging the calcium indicator GCaMP6f under the control of the CaM kinase II promoter-(pENN.AAV.CamKII.GCaMP6f.WPRE.SV40) into the PL (Fig 1D), followed by the implantation of a gradient refractive index (GRIN) lens (Fig 1D-E). The activity of several individual PL neurons in the same field of view was recorded (Fig 1F).

In each behavioral test, we aimed to identify PL neuronal correlates of freezing ^20,21^. At the single cell level during CFC (Fig 1G-I), as well as in the other tasks (Fig S1 D-G), we identified freezing activated (23%), suppressed (47%), or neutral (30%) cells. Freezing activated (FA) and suppressed cells tended to show sharp changes in activity around freezing onset and offset (Fig 1H-I). This data confirms previous findings showing FA cells in the PL ^8,22^. We tracked the same neurons across the different sessions ^23^ (Fig 1J), in pairs of consecutive tasks, such as OF to NO, NO to CFC, and CFC to CFTEST to test their stability (Fig 1K). In NO to CFC, a different set of FA neurons is recruited (Fig 1K), but in OF to NO and CFC to CFTEST, where the context is the same, a significant percentage of cells remained as FA (see methods).

During CFC we detected shock-responding (SR) cells (Fig 1L, Fig S2A) as previously reported ^24^. SR cells showed higher correlations in activity with one another than shock-nonresponding cells (SNR) following FS (Fig 1M, D=0.1554, P<0.0001), even though FS periods were removed from the analysis. SR cells also showed higher correlated activity than SNR cells during CFC before FS occurred (Fig 1O, D=0.1064, P<0.0001), suggesting SR cells might be predetermined to encode the FS. Moreover, SR cells maintained higher correlated activity than SNR cells during CFTEST over the last 2 min (Fig 1N, D=0.1898, P<0.01), the first 2 min (Fig S2D right, D=0.1407, P<0.05) and the entire period (Fig S2D left, D=0.1484, P<0.05). However, this effect was not seen when we tracked SR cells back to the OF (Fig 1P, D=0.1299, P=0.0764; Fig S2B right, D=0.1305, P=0.0741) or NO (Fig 1Q, D=0.1027, P=0.1446; Fig S2C right, D=0.1201, P=0.055) tasks, suggesting that FS event stabilizes the increased correlated activity, that is maintained during recall of the memory trace.

### FS induces a shift in PL neuronal dynamics

To study the response of PL population dynamics to the FS, we conducted a PCA analysis of neuronal activity (Fig 2A), revealing clearly distinct PL clusters between the 2 halves of the CFC (before and after FS, Fig 2B, Fig S2E). This effect was not observed in the other tasks, as evidenced by a higher distance between clusters in the PCA space in CFC related to all other tests (Fig 2C, F_(1.846, 13.54)_=14.74, P<0.001; CFC vs OF, P<0.01 CFC vs NO, P<0.01; CFC vs CFTEST, P<0.05, Tukey’s post-hoc test). To test whether this effect was specific to the PL, we used data obtained from two additional brain areas (Fig 2D-G): one cortical, the Retrosplenial cortex (RSC, N=5 mice) and one subcortical, the Laterodorsal nucleus of the Thalamus (LDn, N=8 mice). We found the centroid distance between the clusters for the 2 halves to be higher for the PL compared to the 2 other regions (Fig 2H, H_(2)_=11.72, P<0.001; PL vs LDn, P<0.05; PL vs RSC, P<0.05, Dunn’s post-hoc test). This finding suggests that PL network activity can be strongly driven by a sudden threat (i.e., FS). These data show that FS during CFC strongly influences population activity within the PL.

**Figure 2.**
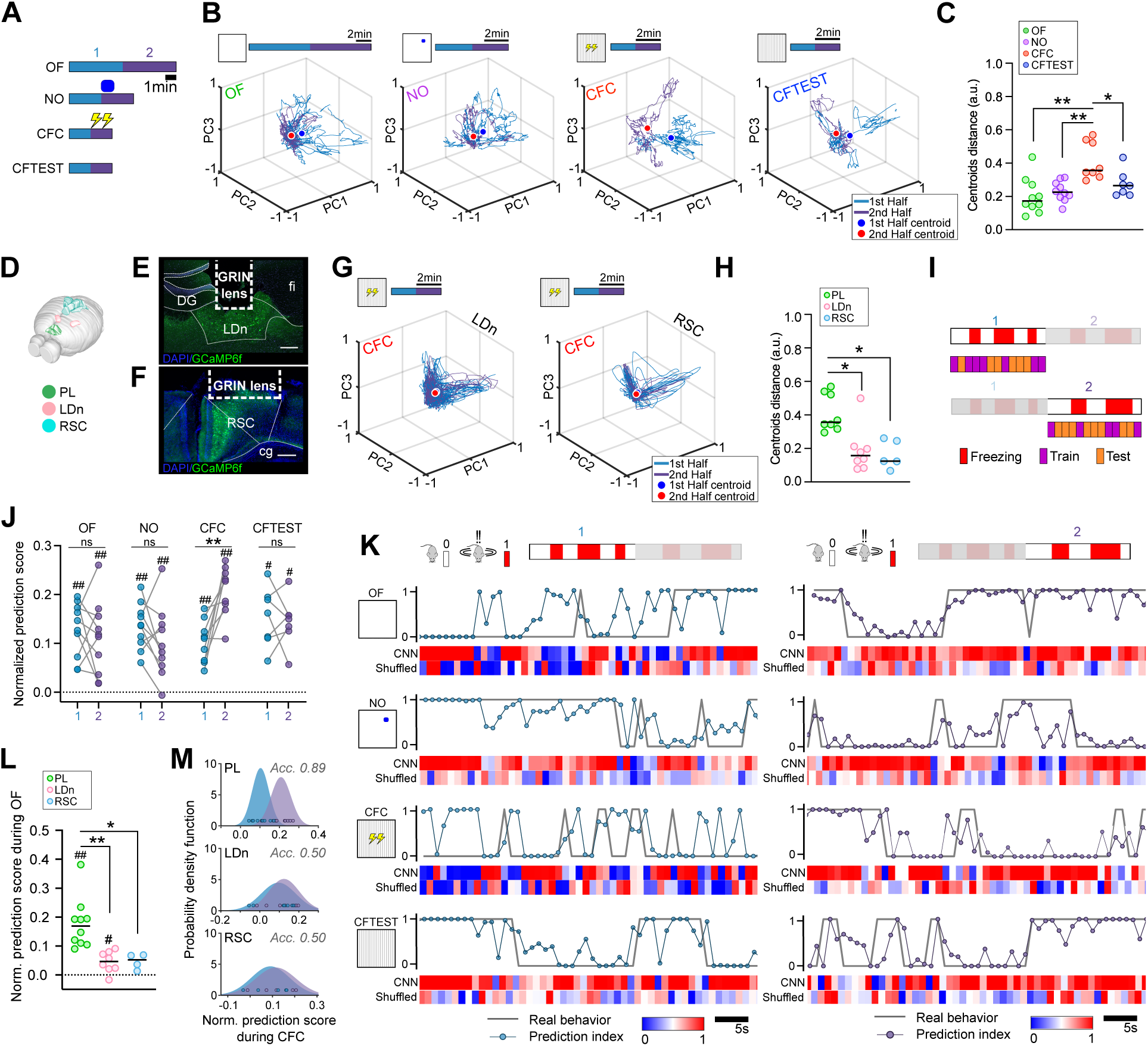
Threat presentation drives a shift in PL, but not in RSC or LDn, neuronal dynamics. **(A)** Schematic showing data splitting (1, 1st half; 2, 2nd half). **(B)** PCA of neuronal activity; color dots represent the cluster centroids. **(C)** Euclidean distances between the centroids of PCA-obtained clusters. **(D)** Brain regions examined with calcium imaging (adapted from the EPFL Blue Brain Atlas). **(E)** Representative viral infection and GRIN lens placement in the LDn and RSC **(F)**. cg, cingulum; DG, Dentate Gyrus; fi, fimbria of the hippocampus. Scale bars 250 µm. **(G)** Representative PCA from the LDn (left) and RSC (right). **(H)** Euclidean distances between centroids of PCA-obtained clusters in PL, LDn, and RSC during CFC. **(I)** Schematic of CNN training and test at 1 (top) and 2 (bottom). **(J)** CNNs-obtained normalized prediction scores distinguishing freezing from non-freezing. **(K)** Examples of prediction scores across time. Top: Real vs predicted freezing. Bottom, performance of CNN with either real or shuffled behavioral annotations. Red blue scale: 1 perfect guess, 0 completely opposite guess. **(L)** CNNs-obtained prediction scores from the PL, LDn, and RSC during OF. **(M)** Naïve Bayes classifier’s probability density functions estimating normalized prediction scores belonging to CFC1 or CFC2, for PL (top), LDn (middle) and RSC (bottom). Acc, Accuracy: fraction of correctly-assigned normalized prediction scores. In relevant panels, black bars represent the median. *P<0.05, **P<0.01, ns = P>0.05, #P<0.05, ##P<0.01.

### Freezing is dynamically encoded in the PL network

Aiming to identify population level representation of freezing, we trained a convolutional neural network (CNN, Fig S3) to classify PL activity as occurring during freezing versus no freezing (Fig S3A). We divided each task into two equal periods, as we did for PCA analysis (see above). We opted for this approach, since, in the CFC and NO tasks, we observed a significant increase in freezing (Fig S1B-C) before and after FS and the novel object introduction, respectively, which we interpreted as a change in behavior towards a threat. Accordingly, we divided and analyzed all tasks in two distinct halves, even if no actual stimulus marked the split (OF and CFTEST) and considered them as two different tasks (Fig 2I). In all tasks (OF 1, OF 2, NO 1, NO 2, CFC 1, CFC 2, CFTEST 1, and CFTEST 2) the CNNs were successfully able to predict freezing from PL neuronal data when compared to their shuffled counterparts (Fig 2J, P<0.05 for all comparisons).

These data suggest that PL neuronal populations achieve distinct functional configurations when the animal is engaged in freezing. Representative time courses of real vs predicted index of freezing are shown in Fig 2K. The same CNN architecture was used with OF data from the RSC and LDn: in both cases, the network performed worse than with PL data (Fig 2L, H_(2)_=15.15, P<0.0001; PL vs LDn, P<0.01; PL vs RSC, P<0.05, Dunn’s test). While in OF, NO, and CFTEST, no differences in the prediction index could be observed between periods 1 and 2 (Fig 2J, OF1 vs OF2, W_(10)_=-13, P=0.5566; NO1 vs NO2, W_(10)_=-27, P=0.1934; CFTEST1 vs CFTEST2, W_(7)_=-4, P=0.8125), a strong increase was observed in CFC2 vs CFC1 (Fig 2J, W_(9)_=43, P<0.01). Although freezing behavior could be predicted by CNNs trained on LDn data (but not for RSC) during OF and CFC (Fig 2L and Fig 3C left, respectively; P<0.05 for all comparisons), we did not observe any change in prediction score following FS during CFC, neither for LDn nor for RSC (Fig 3C left, CFC1 vs CFC2, W_(8)_=18, P=0.25; 3C right, CFC1 vs CFC2, W_(5)_=1, P>0.9999), in contrast to PL data. Moreover, using a naïve Bayes classifier we were able to predict whether a normalized prediction index belongs to CFC1 or CFC2 with 0.89 accuracy for PL, but only with 0.5 for LDn and RSC data (Fig 2M). These data further indicate a network shift specific for PL following FS.

**Figure 3.**
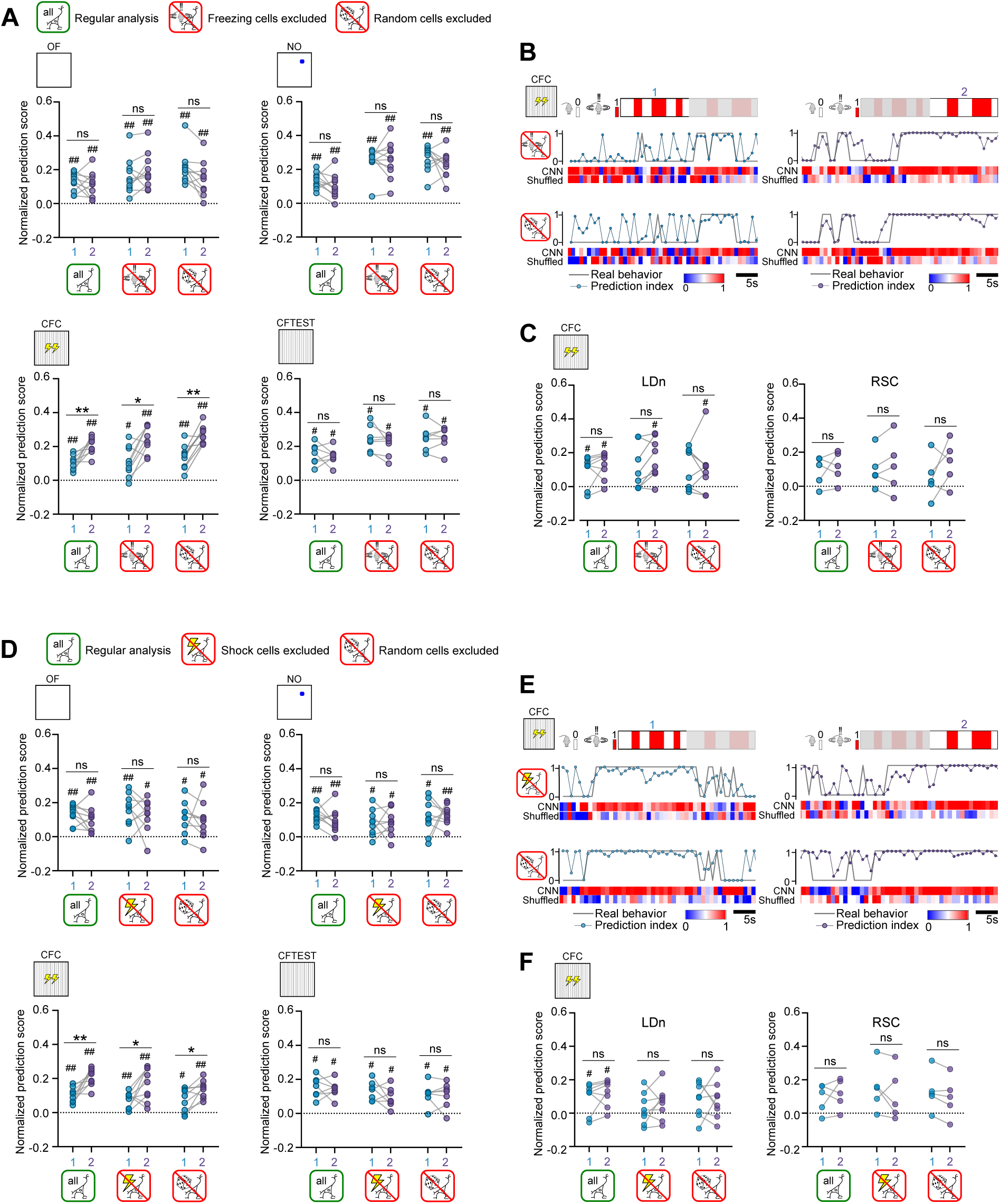
Freezing-activated or shock responding cells do not drive CNN freezing decoding. **(A)** CNNs-obtained freezing prediction scores excluding FA cells during periods 1 vs 2 in the OF (top left), NO (top right), CFC (bottom left), and CFTEST (bottom right) **(B)** Representative freezing prediction score over time in CFC excluding FA cells. **(C)** Freezing prediction scores during CFC from LDn (left) and RSC (right) data excluding FA cells. **(D)** Freezing prediction scores excluding SR cells during periods 1 vs 2 in the OF (top left), NO (top right), CFC (bottom left), and CFTEST (bottom right). **(E)** Representative freezing prediction score over time in CFC excluding SR cells. **(F)** Freezing prediction scores during CFC from LDn (left) and RSC (right) data excluding SR cells. *P<0.05, **P<0.01, ns = P>0.05, #P<0.05, ##P<0.01.

### PL network encoding of freezing is independent of FA or SR cells

We also evaluated whether FA or SR cells played a disproportionate role in the configuration of the PL network (Fig 3). We trained different CNNs, either excluding FA cells, SR cells, or an equivalent randomly drawn population of cells. For all tasks, we found that excluding the FA cells, SR cells or the same number of randomly drawn cells did not affect CNN performances (Fig 3A and 3D, respectively, see Supplementary Table 1 for all comparisons). Representative time courses of real vs predicted freezing during CFC excluding FA or SR cells are shown in Fig 3B and Fig 3E, respectively. Additional examples for other tasks are shown in Fig S4A-B. Normalized prediction score was unaffected by cell exclusion in the LDn and RSC (Fig 3C, Fig 3F). This shows that the CNNs are robust and suggests a populational-level encoding of freezing in the PL network that does not solely rely on the activity of individual cells.

### Threat-driven neuronal dynamics are necessary for fear memory expression

Finally, we tested if shock-state PL associated neuronal activity (Fig 2B) played a role during fear memory expression. We observed subpopulations of neurons that show either decreased or increased activity following FS (Fig 4A). Aiming to verify the contribution of these different subpopulations to memory recall, we applied a novel technology called FLiCRE (fast light and calcium-regulated expression), using an AAV-DJ as a delivery vector, that enables the expression of an opsin whenever a cell is both active and optically-stimulated, with fast temporal resolution ^25^ (Fig 4B). The strategy was to express the inhibitory opsin eNpHR3.0 in neurons that were active, either before or after the FS during CFC (Fig 4A), and silence them during CFTEST (Fig 4C). Neurons in the PL were infected with CaM-uTEVp and neurons tagged expressed the opsin (Fig 4D), with no difference in the percentage of cells infected or tagged between groups (Fig 4E, density of CaM-uTEVp^+^ neurons, U=28, P>0.9999; density of mCherry-eNpHR3.0^+^ neurons, U=22, P=0.5358; % mCherry-eNpHR3.0^+^ / CaM-uTEVp^+^, U=14, P=0.1206). A decrease in freezing upon opto-inhibition with yellow light delivered halfway through recall was observed in mice tagged after (Fig 4F, W_(8)_=-36, P<0.01) but not before FS (Fig 4F, W_(7)_=-8, P=0.5781). Control mice infected with CaM-uTEVp either “non tagged” or “tagged” in the last 2 min of an OF 24 h prior to CFC (Fig S4C), did not show any change in freezing when yellow light was delivered halfway through recall (Fig S4D, W_(5)_=-5, P=0.625 for “non-tagged”; W_(6)_=-15, P=0.1562, for “tagged”). These data show that PL neuronal state following FS is necessary for the proper expression of fear memory.

**Figure 4.**
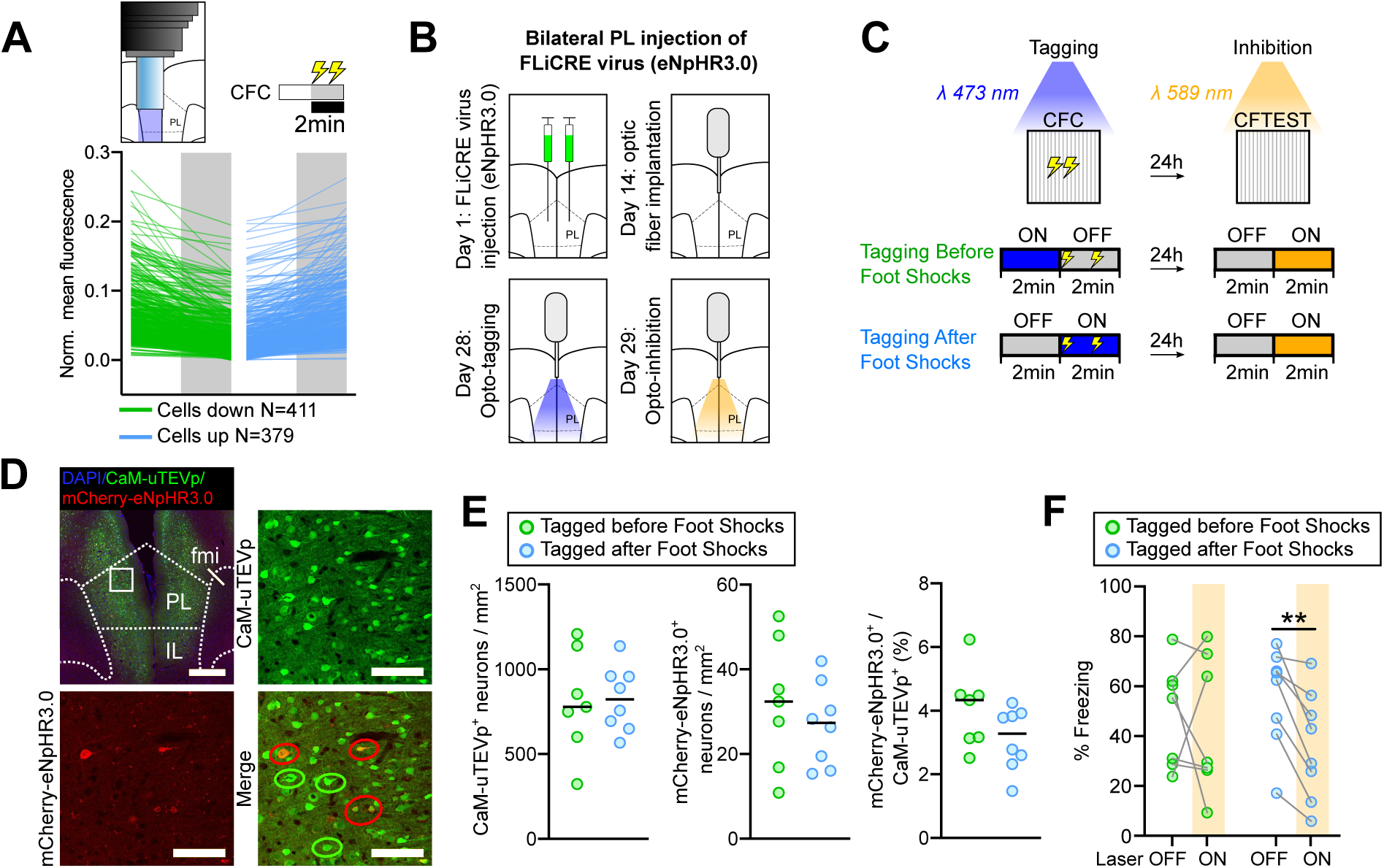
Threat-driven neuronal activity is necessary for proper fear memory expression. **(A)** FS-induced (shaded area) change in activity. **(B)** Timeline of FLiCRE experiment. **(C)** Schematics of experimental setup. **(D)** Example of FLiCRE infection (top left, scale bar 500 μm), magnification showing expression of CaM-uTEVp (top right), mCherry-eNpHR3.0 (bottom left) and merged image (bottom right), showing neurons expressing CaM-uTEVp only (green circles) or both CaM-uTEVp plus mCherry-eNpHR3.0 (red circles). Scale bar 100 μm. fmi, forceps minor of the corpus callosum; IL, Infralimbic cortex. **(E)** Density of CaM-uTEVp^+^ (left) and mCherry-eNpHR3.0^+^ neurons (middle), and percentage of co-expression out of CaM-uTEVp^+^ neurons (right). Each data point corresponds to the mean value of an individual animal. **(F)** Freezing during CFTEST. Black bars in **(E)** represent the median. **P<0.01.

## DISCUSSION

Here we found that freezing is represented at a network level in the PL (and not other regions) at any given time. However, PL freezing representation can shift to internally distinguish between risk-evaluation and threat-dependent fear expression. In fact, following a FS, PL strongly shifts its neuronal state which results in an enhancement of freezing representation. This shift in PL activity is necessary for a proper memory-related fear representation the day after. Importantly, FS had no effect on network perturbation when tested in control regions (here RSC and LDn), and freezing could not always be decoded from these regions’ activity. These data show the involvement of the PL in processing fear-related behavior at the single cell (presence of FA and SR cells) and population level activity (as rapid shifts in neuronal state), revealing the unique role of PL in adjusting its activity towards threats to better represent and recall fear.

The PL cortex is a key region for both risk assessment and fear representation ^8,22,26^. The use of 3 behavioral tasks (OF, NO, CFC) enabled us to study the dynamics of PL single neurons, populations, and neuronal networks underlying fear. We found FA cells in every task. These cells are unstable since new subpopulations of cells are recruited in different tasks. However, when mice are placed back in the same context (NO after OF or CFTEST after CFC) the percentage of overlapping between FA cells increases over chance, suggesting spatial memory features despite their instability. On the other hand, their instability may reflect PL capability to evaluate the present situation and adjust its neuronal representation accordingly. Drawing a parallel with the amygdala, neurons in this area have a more stable encoding of exploratory and defensive behaviors across different tasks ^27^. These findings are in line with the proposed role of the amygdala as a direct regulatory center for emotional behaviors, and the mPFC involved in the evaluation and organization of behavior.

Then we analyzed PL SR cells and their engagement in fear memory recall. SR cells showed higher correlated activity during CFC, creating an ensemble of cells that maintained their correlation 1 day after, presenting an early 1-day memory signature. During CFC, the higher correlated activity of SR cells was detected even before FS was delivered, which might suggest these cells were pre-allocated, consistent with the current theory of CREB-allocation of memory-related neuronal ensembles in other brain areas ^28^. This feature was not observed when SR cells were tracked back to NO and OF, suggesting that the pre-allocation occurs during the present task (CFC) and not before. The mPFC has been largely studied for its role in remote memory recall ^29–31^. mPFC cells that are active during the acquisition of fear memories are tagged early, via epigenetic and synaptic modifications ^32,33^, and undergo synaptic rearrangement immediately after training ^34^. mPFC reactivation is necessary for fear memory recall at remote time points ^31,34,35^, indicating its crucial role in long-term memory stabilization. Kitamura et al. ^24^ also reported the presence of SR cells in the mPFC, showing that they are preferentially reactivated at a remote time point (15 days after training), but not in a 1-day-after test, and suggested they may constitute the mPFC remote memory engram. To our knowledge, no evidence pointed to a different recruitment of mPFC SR cells as early as after 1 day. However, it has been proposed that a possible way the prefrontal cortex increases stability is via the formation of ensembles of cells that are synchronously activated ^36^, stabilizing neuronal activity and allowing memory recall via pattern completion ^37^. Our results show the presence of synchronized ensembles both during training but also 1 day after, and are in line with Zelikowsky et al. ^38^ findings, showing mPFC representation of both contextual and emotional aspects of memory.

Using a deep learning approach, we were able to decode freezing from PL network activity in all tasks with higher accuracy than in other brain regions (RSC and LDn tested here). This implies that there are freezing signatures that can be detected by the PL network. The capacity of the network to predict freezing is preserved even after the removal of either random, FA or SR cells (Fig 3). These data indicate that networks are robust and that all cells, independent of its preferential activity, carry freezing-related information. However, SR cells might be driving the FS-dependent neuronal shift detected at a population level (Fig 2). The FS-induced change in PL neuronal configuration improved freezing decoding (Fig 2J). This is in line with a study showing a shift in the dorsomedial prefrontal cortex (dmPFC) population neural state during conditioned avoidance ^14^. However, while Jercog et al. ^14^ identified a neuronal dissociation between cue encoding versus escape in the dmPFC, we show that PL neuronal population can predict freezing in different contexts. Importantly, our data showed 2 PL dynamics states underlying freezing, each corresponding to specific internal states occurring before and after FS. Notably, there were no observed changes in the neural population state or decoding of freezing behavior in the RSC or the LDn following FS. These data support the hypothesis that PL is involved in risk evaluation, as fear is differentially represented in the PL in the face of a threat.

Finally, we found evidence for the functional importance of FS-driven PL state shift: we used a novel approach called FLiCRE ^25^ to “tag” different subpopulations of cells active either before or after FS presentation during CFC. Neurons expressing eNpHR3.0 were inactivated optogenetically halfway through CFTEST. Mice whose PL neurons were “tagged” after, but not before FS, showed impaired memory performances (i.e., decreased freezing levels) upon inhibition. This result pointed to the importance of the PL neuronal shift in controlling fear expression linked to previously learned fear experiences.

In conclusion, this study supports and brings new evidence for a neuronal organization within the PL that represents fear. At a single cell level, selected PL neurons are preferentially more active during freezing and FS, and they can retain this information in a context-dependent manner as early as 1 day after acquisition. At a population level, PL neural representation accompanying freezing can be better decoded than in other brain areas, is strongly improved in the face of a real threat and is necessary for proper fear memory expression, consistent with a role for the PL in risk evaluation.

## ACKNOWLEDGMENTS

We thank Colleen J. Gillon for the helpful discussions, comments, and revisions. We also would like to thank Bianca Silva and Gabrielle Girardeau for commenting on a previous version of the paper. This work was supported by grants to G.V. from ANR (ANR-20-CE16-0015-01), Emergence(s) Ville de Paris (2019 DAE 81), and a NARSAD Young Investigator Grant from the Brain and Behavior Research Foundation (#29489). J.P.C. was partially funded by a FONDECYT (N**°**3170497) grant and a MSCA postdoctoral fellowship (#896825).

## AUTHOR CONTRIBUTION

JPC, CP, IA, and GV designed the experiments. JPC conducted all optogenetics and calcium imaging experiments. JPC, CP, NT and IA conducted data analysis. MB prepared the viruses for FLiCRE experiments. GV, JPC, and CP wrote the paper.

## METHODS

### RESOURCE AVAILABILITY

#### Lead contact

Further information and requests should be directed to and will be fulfilled by the lead contact, Gisella Vetere.

#### Materials availability

This study did not generate new unique materials.

### EXPERIMENTAL MODEL AND SUBJECT DETAILS

#### Animals

129/Sv x C57BL adult (7-8 weeks) male and female mice were used. They were kept in groups of 2 to 4 during the whole protocol, under a standard lab diet with food and water *ad-libitum*, on a 12 h light/dark cycle. Experiments were done typically between 9:00 and 14:00 h, during the light phase. All procedures were performed in accordance with the official European guidelines for the care and use of laboratory animals (86/609/EEC), in accordance with the Policies of the French Committee of Ethics (Decrees n° 87– 848 and n° 2001–464) and after approval by ethical committee (reference: 2019-22).

### METHOD DETAILS

#### Surgical procedures

For every surgery, mice were anesthetized with an i.p. injection of a mixture of ketamine/xylazine 100/10 mg/kg. For calcium imaging experiments, 1 µL of a virus encoding AAV5.CamKIIa.GCaMP6f (pENN.AAV.CamKII.GCaMP6f.WPRE.SV40, Addgene viral prep 100834-AAV5) virus was injected unilaterally in the PL (AP +1.9, ML -0.25, DV -2.0 ^39^) using a glass micropipette. Two weeks later, mice were implanted with a GRIN lens (GO!Foton, 1mm diameter, 4 mm length) which was slowly lowered right to the viral injection site. To help make a track for the GRIN lens, a 1 mm diameter needle was stereotaxically inserted and then removed from the brain. Two anchor screws (FST) were used to add more grip for the cement (Lang Dental). The surface of the lens was protected with a small piece of parafilm and kwik-cast silicone (WPI). Following two weeks, a magnetic base plate for the miniscope was implanted, and imaging started the next day. Two additional cohorts of mice were used to image the right LDn (N=8) and the RSC (N=5) following the same procedures mentioned above, except the viral injection and GRIN lens (0.5mm for LDn and 1mm for RSC) were both aimed at AP -1.2, ML -1.2, DV -2.6 for the LDn; and AP -1.5, ML -0.3, DV -1.0 for the RSC mice ^39^. Prior to behavioral experiments, mice were handled and habituated to the miniscope for around 5 min, for at least 3 consecutive days. For the optogenetic tagging of active neurons we bilaterally injected 0.6 µL of 1:1:1 mix of AAVDJ-GFP-CaM-uTEVp, AAVDJ-Nrxn3b-Nav1.6-MKII-f-hLOV1-TEVcs(ENL YQ/M)-tTA-VP16 and AAVDJ-TRE:mCherry-p2a-eNpHR3.0 virus aimed at the PL cortex (AP +1.9, ML - 0.25, DV -2.0 ^39^). Following two weeks mice were implanted with a single 200 µm optic fiber (0.37 NA) coupled to a ceramic ferrule connector aimed to the midline above the PL to cover both hemispheres (AP +1.9, ML 0.0, DV -1.7). Fibers extending around 1.9 mm beyond ferrules, were anchored to the skull with black dental cement and 2 screws. Mice were then allowed to recover for 1 week and were habituated to both experimenter and optic fiber coupling for 3 consecutive days before experiments.

#### Viral Production

FLiCRE expression vectors (addgene) were packaged into AAV serotype DJ (cell biolabs #VPK-400-DJ). at St. Jude Children’s Research Hospital. Adenoviral helper genes were provided using the plasmid pHGTI-adeno1 ^40^. Plasmids were transfected into Adherent 293T cells (ATCC CRL-3216), 10x15 cm dishes per construct, at a 1:1:1 ratio using PEI-polyethyleneimine “max,” (Polysciences #24765). Media was changed to serum free DMEM at 16 hours post transfection. Cell supernatants and pellets were harvested 72 hours post transfection. Cell pellets were lysed by 5 freeze-thaw cycles, lysate was collected and diluted in serum free DMEM to a volume of 40 mL and pegylated with 40% PEG8000 (Fisher) at a 1:5 overall volume (10 mL PEG). Supernatants were directly pegylated with 40% PEG8000 at a 1:5 overall volume. Pegylated lysates and supernatants were incubated at 4°C for two hours and then centrifuged for 30 minutes at 4°C (4000 g). The PEG containing pellets were resuspended in 10 mM Tris, 10 mM Bis-Tris-Propane (pH 9). The resuspended sample was treated with benzonase (Sigma) and loaded onto a two-step CsCl step gradient ^41^ in a thick wall ultracentrifuge tube (Beckman Cat# 360743) Samples were loaded onto an ultracentrifuge (Sw32Ti rotor) and spun at 24,600 rpm at room temperature for 20 hours. The full particle containing fraction was isolated, loaded onto a dialysis cassette (Thermo cat# 66810) and dialysed 3x in 1xPBS. Dialyzed virus was collected, filtered through a 0.2 μM filter and concentrated by a 100 Kda filter (Amicon) to a volume of 0.5 mL. AAV vectors were titered by qPCR using serial dilutions of purified virus compared against linearized plasmid standard references. Viruses were aliquoted and stored at -80°C until use.

#### Behavioral procedures

##### Open field (OF)

Mice were connected to the miniscope and were placed in a familiar environment while acquisition parameters were adjusted. Then, they were placed in the center of the OF (47x47x35 cm arena) while their behavior was recorded by a webcam placed on top, and they were allowed to explore for 10 min.

##### Novel object test (NO)

Two days after the OF test, mice were returned to the same arena. Following 3 min of free exploration, an object (a small 5.5x4.5x3.0 cm cardboard box) that was completely new to them was placed in the opposite corner to the one the mouse was closest to, around 10 cm from each wall. Mice were allowed to explore for an additional 3 min period.

##### Contextual fear conditioning (CFC) and recall (CFTEST)

Two days after the NO test, mice were placed in an enclosed fear conditioning cage (IMETRONIC, France). After 2 min, mice were subjected to two FS (0.2 mA, 2 s), spaced 1 min apart. The whole training took 4 min. The next day (CFTEST), mice were placed back in the same environment for 4 min, while no FS were delivered.

#### Mice position tracking and behavioral scoring

Mouse position was determined by detecting the centroid of the mouse body shape in the field using the Bonsai software ^42^. Freezing was scored semi-automatically using Bonsai: first, we ran a workflow to detect periods of immobility, then a trained experimenter ran a frame-by-frame analysis of the detected periods of immobility. The mice from the LDn and RSC cohorts were used for a different study, but prior to any other experiment being run, they were subjected to the OF and then to CFC under the same conditions as the PL-implanted mice.

#### In vivo calcium Imaging

We visualized the activity of an average of 102.5 cells per session (range 71 to 128 cells per session per mouse; OF and NO, N=10 mice; CFC, N=9 mice; CFTEST, N=7 mice). Ca^2+^ traces were obtained using min1pipe ^43^ running in MATLAB 2021a. To classify neurons as either freezing activated, suppressed, or neutral, the mean activity of each neuron during freezing vs no freezing episodes was compared (one tail t-test, P<0.01). Overlapping freezing activated cells were calculated by counting the number of cells that were considered as such in a pair of consecutive tasks (i.e. OF vs NO, NO vs CFC, and CFC vs CFTEST). Then we randomly selected pairs of cells from two consecutive tasks 5000 times to obtain a distribution of overlapping freezing-activated cells. If the real number of overlapping cells was above the 95th percentile of the null distribution, we considered the overlap significant. Changes in PL population activity during each task were analyzed using PCA with frames (time) as observations and neurons as variables. In each task, the whole period was split into two halves, which in the case of NO and CFC are divided by the introduction of the object and the first FS, respectively. The Euclidean distance between the two cluster centroids was used as an index of how different the population activity was.

#### Cell registration across tasks

The tracking of the same cells active over different tasks (different days) was conducted with CellReg ^44^, a probabilistic approach that allows cell registration with minimal user intervention. Briefly, we first obtained the neuronal spatial footprint from each imaging session with Min1pipe and used one of them as a reference. The alignment of two sessions was attempted by either rotations, translations, or non-rigid corrections to the footprints, with the latter being the most frequently used. Final registration was computed by spatial correlations or centroid distances of potential cell pairs.

#### SR cell detection

To detect SR cells, cell activity was examined 4 seconds before and 4 seconds after shock onset. For each cell and each shock, the 8 seconds of activity were z-scored using the median and standard deviation of the first 4 seconds. The resulting post-onset activity was averaged between both shocks, giving us the ‘alpha score’ (i.e., the average activity of the 4 seconds after first shock onset and 4 seconds after second shock onset, each normalized to the 4 seconds before the corresponding shock onset). “Random” alpha scores were obtained for each cell by shuffling the position of the shock onsets 1000 times.

#### Correlations of calcium traces

We analyzed whether any pair of cells are generally active at the same time. To do so, for each animal and in each experiment, we used Matlab’s built-in cross-correlation function ‘xcorr’ to correlate the normalized calcium traces of each pair of cells. For each cell pair, the correlation value used is the maximum of all possible normalized cross-correlations of the pair with a maximum shift of +- 40 frames (+- 2 seconds), thus obtaining the correlation *r_xy_* of neurons x and y:

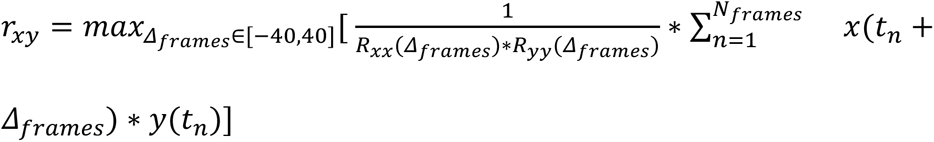

Where x is the calcium trace of a neuron, y the calcium trace of another neuron, and *R_xx_* and *R_yy_* autocorrelations of said neurons.

#### Optogenetic tagging and inhibition of foot shock-state associated cells

Mice were subjected to a 4 min OF, followed by CFC and CFTEST under the same conditions as for calcium imaging experiments. During CFC, PL was constantly illuminated with blue light laser (473 nm, 10 mW) either before (first 2 min) or after FS (last 2 min). The following day mice were re-exposed to the same context and PL was constantly illuminated with yellow light laser (589 nm, 5 mW) during the last 2 min of CFTEST.

#### Convolutional Neural Network analysis

We designed a convolutional deep neural network (CNN) to classify neural data (calcium traces) into two possible categories (observed behavioral states), i.e., freezing and no freezing. We concluded that if the network was able to successfully differentiate between two behaviors, then these must be encoded differently in the PL at a population level. Neural data was formatted into one-second-long bouts of z-scored calcium traces from one experiment. Only the 60 most active neurons (average z-scored fluorescence over the whole experiment) were considered for each mouse in each experiment in order to pool the data across mice for training the network. We verified whether this arbitrary threshold had an effect on CNN results by correlating normalized prediction scores with the corresponding threshold values of activity for selecting cells for each animal in each experiment. No effect was detected in any of the experiments (Pearson correlations, OF1: p=0.29, R²=0.14, OF2: p=0.37, R²=0.10, NO1: p=0.63, R²=0.03, NO2: p=0.25, R²=0.16, CFC1: p=0.87, R²=0.004, CFC2: p=0.70, R²=0.02, CFTEST1: p=0.59, R²=0.06, CFTEST2: p=0.16, R²=0.36). These episodes were labeled as freezing (1) or no freezing (0). The input data for each trial, i.e. a 1-second bout of calcium traces for 60 neurons was then passed through two convolutional layers, followed by a series of fully connected layers. A rectified linear (ReLU) non-linearity was applied to each layer. The output layer, comprising 2 units, was transformed using a softmax in order to obtain probabilities, or prediction indices, for each class. See CNN structure below; and Fig S3. The network was trained on this binary classification task with a negative log-likelihood loss, and using backpropagation with Adam optimization. The prediction index reported in the paper is the probability assigned by the network to the “freezing” class. For each experiment, the data was split between training and testing. Training data consisted in equal parts of freezing and no freezing bouts. The size of the training data was determined so that every CNN was trained on the same amount of data. To do so, the train/test split was done on every experiment by 1) withholding a third of the data for testing and 2) random subsampling of trials to obtain a 50%/50% freezing versus no freezing trials split in the training data. Random subsampling was repeated to match the size of the experiment with the smallest training set. As a result, CNNs trained on the different experiments were exposed to the exact same amount of training data while still being trained on a 50%/50% split between categories. We trained multiple CNNs and retained the best-scoring network to predict the labels for the test data. We scored the results by running the network separately 100 times, on 100 different train/test splits: this allowed us to have each bout of activity appear in the test data in multiple runs (about 20 times). Some bouts did not reach a minimum of 20 individual guesses, and were then not considered for analysis (hence the slight difference in some displayed behavior bouts, see Fig 3B, 3E and S4). By averaging those repeated prediction indices, we obtained a robust prediction index for each bout, displayed as dots in Fig 2K, Fig 3B, Fig 3E, Fig S4A, Fig S4B. These robust prediction indices were then individually compared to the real behavior (see the ‘CNN’ colored lines in Fig 2K, Fig 3B, Fig 3E, Fig 3K, Fig S4A, Fig S4B), grouped by animal, and averaged, with equal weight for freezing and no freezing trials:

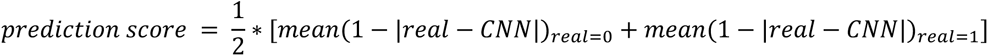

This allowed us to avoid a bias caused by an unequal number of freezing and no freezing trials for individual animals or experiments. We obtained prediction scores for each experiment and each animal using the correct annotation and then repeated this process 20 times with shuffled annotations (see the ‘Shuffled’ colored lines in Fig 2K, Fig 3E, Fig 3K, Fig S4A, Fig S4B to obtain chance prediction scores, which, when subtracted from the real prediction scores, gave the normalized prediction scores reported in Fig 2J-M and Fig 3. For the exclusion training (Fig 3 and Fig S4), CNNs were trained the exact same way, but excluding either the activity of freezing-activated cells or SR cells when identifying the 60 most active cells. As a control, CNNs were trained similarly, by excluding the same number of random cells from the 60 most active cells identified.

The CNN structure was as follows:

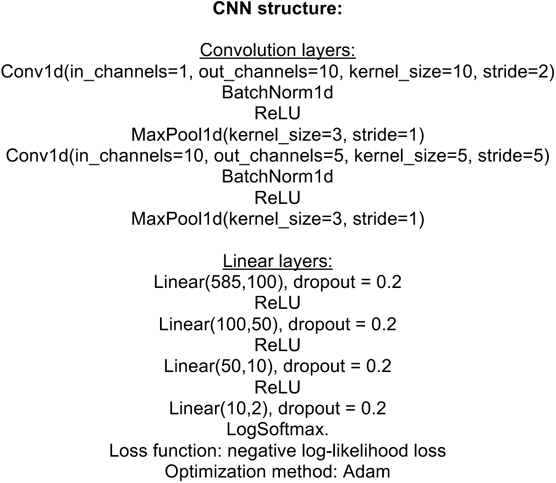

We also fitted a naïve Bayes classifier to normalized prediction scores of CFC1 and CFC2 (2 data points per animal), and obtained the corresponding fitted probability density functions. Thus, we could then estimate, for each data point, its probability of being before (CFC1) or after (CFC2) the shocks and obtain an accuracy for PL, LDn, and RSC data (Fig 2M).

#### Histology

After completion of the behavioral battery, mice were deeply anesthetized and transcardially perfused with around 50 mL of saline and then 50 mL of 4% PFA. The brains were extracted and kept in PFA for 24 h and then placed in 30% sucrose until they sank. Brains were then sliced with a cryostat at 50 µm. Slices were kept in 0.02% azide PBS, stained with DAPI (1:10000), and mounted with PermaFluor (Thermo Scientific) for subsequent histological analysis. QuPath v.0.4.0 ^45^ was used to analyze FLiCRE infection and tagging within the manually selected PL in both hemispheres. For each animal two to three slices were evaluated. During image processing and analysis, the experimenter remained blind to the conditions. Detection and quantification of CaM-uTEVp^+^ cells was accomplished through a semi-automatic approach using the “Cell detection” plugin in QuPath. To assess tagging within infected cells, mCherry-eNpHR3.0^+^ cells were identified exclusively within CaM-uTEVp^+^ detected cells. For comparing the tagging before and after treatment, the percentage of mCherry-eNpHR3.0^+^ to CaM-uTEVp^+^ cells was calculated for each slice and averaged for the respective animal. The absolute amount of mCherry-eNpHR3.0^+^ and CaM-uTEVp^+^ cells was normalized based on the measured area and reported per 1 mm^2^. Representative images were created with ImageJ-win64 and Inkscape.

#### Statistical analysis

The sample size was defined by what is already published in similar studies and not using statistical approaches. Data were plotted and analyzed with GraphPad Prism 8.0.1 software or custom scripts written in MATLAB or Python. For analyses aimed at comparing two groups, we used non-parametric (two-tailed) paired or unpaired tests, as applicable. For analyses across tasks, we used a Mixed effects model test (as implemented in GraphPad Prism 8.0.1) followed by Tukey’s post hoc test for multiple comparisons. Mixed effects model tests can handle missing values, allowing the results to be interpreted like a repeated measures ANOVA. Differences in frequency distributions were assessed with the Kolmogorov-Smirnov test. Further details about statistical tests can be found in Supplementary Table 1.

## REFERENCES

1. Anastasiades, P.G., and Carter, A.G. (2021). Circuit organization of the rodent medial prefrontal cortex. Trends Neurosci 44, 550–563. 10.1016/j.tins.2021.03.006.

2. Orsini, C.A., Kim, J.H., Knapska, E., and Maren, S. (2011). Hippocampal and Prefrontal Projections to the Basal Amygdala Mediate Contextual Regulation of Fear after Extinction. J Neurosci 31, 17269–17277. 10.1523/jneurosci.4095-11.2011.

3. Salzman, C.D., and Fusi, S. (2010). Emotion, Cognition, and Mental State Representation in Amygdala and Prefrontal Cortex. Annual Review of Neuroscience 33, 173–202. 10.1146/annurev.neuro.051508.135256.

4. Do-Monte, F.H., Quiñones-Laracuente, K., and Quirk, G.J. (2015). A temporal shift in the circuits mediating retrieval of fear memory. Nature 519, 460–463. 10.1038/nature14030.

5. Corcoran, K.A., and Quirk, G.J. (2007). Activity in Prelimbic Cortex Is Necessary for the Expression of Learned, But Not Innate, Fears. J Neurosci 27, 840–844. 10.1523/jneurosci.5327-06.2007.

6. Stern, C.A.J., Monte, F.H.M.D., Gazarini, L., Carobrez, A.P., and Bertoglio, L.J. (2010). Activity in prelimbic cortex is required for adjusting the anxiety response level during the elevated plus-maze retest. Neuroscience 170, 214–222. 10.1016/j.neuroscience.2010.06.080.

7. Sierra-Mercado, D., Padilla-Coreano, N., and Quirk, G.J. (2011). Dissociable Roles of Prelimbic and Infralimbic Cortices, Ventral Hippocampus, and Basolateral Amygdala in the Expression and Extinction of Conditioned Fear. Neuropsychopharmacol 36, 529–538. 10.1038/npp.2010.184.

8. Burgos-Robles, A., Vidal-Gonzalez, I., and Quirk, G.J. (2009). Sustained Conditioned Responses in Prelimbic Prefrontal Neurons Are Correlated with Fear Expression and Extinction Failure. J Neurosci 29, 8474–8482. 10.1523/jneurosci.0378-09.2009.

9. Quiñones-Laracuente, K., Vega-Medina, A., and Quirk, G.J. (2021). Time-Dependent Recruitment of Prelimbic Prefrontal Circuits for Retrieval of Fear Memory. Front Behav Neurosci 15, 665116. 10.3389/fnbeh.2021.665116.

10. Jinks, A.L., and McGregor, I.S. (1997). Modulation of anxiety-related behaviours following lesions of the prelimbic or infralimbic cortex in the rat. 1–10.

11. Dejean, C., Courtin, J., Karalis, N., Chaudun, F., Wurtz, H., Bienvenu, T.C.M., and Herry, C. (2016). Prefrontal neuronal assemblies temporally control fear behaviour. Nature 535, 420–424. 10.1038/nature18630.

12. Suzuki, S., Saitoh, A., Ohashi, M., Yamada, M., Oka, J.-I., and Yamada, M. (2016). The infralimbic and prelimbic medial prefrontal cortices have differential functions in the expression of anxiety-like behaviors in mice. Behavioural Brain Research 304, 120–124. 10.1016/j.bbr.2016.01.044.

13. Armbruster, D.J.N., Ueltzhöffer, K., Basten, U., and Fiebach, C.J. (2012). Prefrontal Cortical Mechanisms Underlying Individual Differences in Cognitive Flexibility and Stability. J Cognitive Neurosci 24, 2385–2399. 10.1162/jocn_a_00286.

14. Jercog, D., Winke, N., Sung, K., Fernandez, M.M., Francioni, C., Rajot, D., Courtin, J., Chaudun, F., Jercog, P.E., Valerio, S., et al. (2021). Dynamical prefrontal population coding during defensive behaviours. Nature 595, 690–694. 10.1038/s41586-021-03726-6.

15. Blanchard, D.C., and Blanchard, R.J. (2008). Chapter 2.4 Defensive behaviors, fear, and anxiety. Handb. Behav. Neurosci. 17, 63–79. 10.1016/s1569-7339(07)00005-7.

16. Roy, V., and Chapillon, P. (2004). Further evidences that risk assessment and object exploration behaviours are useful to evaluate emotional reactivity in rodents. Behav Brain Res 154, 439–448. 10.1016/j.bbr.2004.03.010.

17. Lezak, K.R., Missig, G., and Jr, W.A.C. (2017). Behavioral methods to study anxiety in rodents. Dialogues Clin. Neurosci. 19, 181–191. 10.31887/dcns.2017.19.2/wcarlezon.

18. Aharoni, D., and Hoogland, T.M. (2019). Circuit Investigations With Open-Source Miniaturized Microscopes: Past, Present and Future. Frontiers in Cellular Neuroscience 13, e1701548–12. 10.3389/fncel.2019.00141.

19. Ghosh, K.K., Burns, L.D., Cocker, E.D., Nimmerjahn, A., Ziv, Y., Gamal, A.E., and Schnitzer, M.J. (2011). Miniaturized integration of a fluorescence microscope. Nat Methods 8, 871–878. 10.1038/nmeth.1694.

20. Adhikari, A., Topiwala, M.A., and Gordon, J.A. (2011). Single Units in the Medial Prefrontal Cortex with Anxiety-Related Firing Patterns Are Preferentially Influenced by Ventral Hippocampal Activity. Neuron 71, 898–910. 10.1016/j.neuron.2011.07.027.

21. Adhikari, A., Topiwala, M.A., and Gordon, J.A. (2010). Synchronized Activity between the Ventral Hippocampus and the Medial Prefrontal Cortex during Anxiety. Neuron 65, 257–269. 10.1016/j.neuron.2009.12.002.

22. Bagur, S., Lefort, J.M., Lacroix, M.M., Lavilléon, G. de, Herry, C., Chouvaeff, M., Billand, C., Geoffroy, H., and Benchenane, K. (2021). Breathing-driven prefrontal oscillations regulate maintenance of conditioned-fear evoked freezing independently of initiation. Nat Commun 12, 2605. 10.1038/s41467-021-22798-6.

23. Sheintuch, L., Rubin, A., Brande-Eilat, N., Geva, N., Sadeh, N., Pinchasof, O., and Ziv, Y. (2017). Tracking the Same Neurons across Multiple Days in Ca2+ Imaging Data. Cell Reports 21, 1102–1115. 10.1016/j.celrep.2017.10.013.

24. Kitamura, T., Ogawa, S.K., Roy, D.S., Okuyama, T., Morrissey, M.D., Smith, L.M., Redondo, R.L., and Tonegawa, S. (2017). Engrams and circuits crucial for systems consolidation of a memory. Science 356, 73–78. 10.1126/science.aam6808.

25. Kim, C.K., Sanchez, M.I., Hoerbelt, P., Fenno, L.E., Malenka, R.C., Deisseroth, K., and Ting, A.Y. (2020). A Molecular Calcium Integrator Reveals a Striatal Cell Type Driving Aversion. Cell 183, 2003–2019.e16. 10.1016/j.cell.2020.11.015.

26. Giustino, T.F., and Maren, S. (2015). The Role of the Medial Prefrontal Cortex in the Conditioning and Extinction of Fear. Front Behav Neurosci 9, 298. 10.3389/fnbeh.2015.00298.

27. Gründemann, J., Bitterman, Y., Lu, T., Krabbe, S., Grewe, B.F., Schnitzer, M.J., and Lüthi, A. (2019). Amygdala ensembles encode behavioral states. Science 364, eaav8736–11. 10.1126/science.aav8736.

28. Zhou, Y., Won, J., Karlsson, M.G., Zhou, M., Rogerson, T., Balaji, J., Neve, R., Poirazi, P., and Silva, A.J. (2009). CREB regulates excitability and the allocation of memory to subsets of neurons in the amygdala. Nat Neurosci 12, 1438–1443. 10.1038/nn.2405.

29. Squire, L.R., and Alvarez, P. (1995). Retrograde amnesia and memory consolidation: a neurobiological perspective. Curr Opin Neurobiol 5, 169–177. 10.1016/0959-4388(95)80023-9.

30. Wiltgen, B.J., Brown, R.A.M., Talton, L.E., and Silva, A.J. (2004). New Circuits for Old Memories The Role of the Neocortex in Consolidation. Neuron 44, 101–108. 10.1016/j.neuron.2004.09.015.

31. Frankland, P.W., and Bontempi, B. (2005). The organization of recent and remote memories. Nature Reviews Neuroscience 6, 119–130. 10.1038/nrn1607.

32. Lesburguères, E., Gobbo, O.L., Alaux-Cantin, S., Hambucken, A., Trifilieff, P., and Bontempi, B. (2011). Early Tagging of Cortical Networks Is Required for the Formation of Enduring Associative Memory. Science 331, 924–928. 10.1126/science.1196164.

33. Vetere, G., Borreca, A., Pignataro, A., Conforto, G., Giustizieri, M., Marinelli, S., and Ammassari-Teule, M. (2019). Coincident Pre- and Post-Synaptic Cortical Remodelling Disengages Episodic Memory from Its Original Context. 1–11. 10.1007/s12035-019-01652-3.

34. Vetere, G., Restivo, L., and J. C. cristina (2011). Spine growth in the anterior cingulate cortex is necessary for the consolidation of contextual fear memory. PNAS, 1–5. 10.1073/pnas.1016275108/-/dcsupplemental/pnas.201016275si.pdf.

35. Restivo, L., Vetere, G., Bontempi, B., and Ammassari-Teule, M. (2009). The formation of recent and remote memory is associated with time-dependent formation of dendritic spines in the hippocampus and anterior cingulate cortex. The Journal of neuroscience : the official journal of the Society for Neuroscience 29, 8206–8214. 10.1523/jneurosci.0966-09.2009.

36. Hanganu-Opatz, I.L., Klausberger, T., Sigurdsson, T., Nieder, A., Jacob, S.N., Bartos, M., Sauer, J.-F., Durstewitz, D., Leibold, C., and Diester, I. (2023). Resolving the prefrontal mechanisms of adaptive cognitive behaviors: A cross-species perspective. Neuron 111, 1020–1036. 10.1016/j.neuron.2023.03.017.

37. Carrillo-Reid, L., and Yuste, R. (2020). Playing the piano with the cortex: role of neuronal ensembles and pattern completion in perception and behavior. Curr Opin Neurobiol 64, 89–95. 10.1016/j.conb.2020.03.014.

38. Zelikowsky, M., Hersman, S., Chawla, M.K., Barnes, C.A., and Fanselow, M.S. (2014). Neuronal Ensembles in Amygdala, Hippocampus, and Prefrontal Cortex Track Differential Components of Contextual Fear. J Neurosci 34, 8462–8466. 10.1523/jneurosci.3624-13.2014.

39. 39. Paxinos, G., and Franklin, K.B.J. (2004). The Mouse Brain in Stereotaxic Coordinates Elsevier, ed.

40. Streck, C.J., Dickson, P.V., Ng, C.Y.C., Zhou, J., Gray, J.T., Nathwani, A.C., and Davidoff, A.M. (2005). Adeno-Associated Virus Vector-Mediated Systemic Delivery of IFN-β Combined with Low-Dose Cyclophosphamide Affects Tumor Regression in Murine Neuroblastoma Models. 11, 6020–6029. 10.1158/1078-0432.ccr-05-0502.

41. Gray, S.J., Choi, V.W., Asokan, A., Haberman, R.A., McCown, T.J., and Samulski, R.J. (2011). Production of recombinant adeno-associated viral vectors and use in in vitro and in vivo administration. Curr Protoc Neurosci Chapter 4, Unit 4.17. 10.1002/0471142301.ns0417s57.

42. Lopes, G., Bonacchi, N., Frazão, J., Neto, J.P., Atallah, B.V., Soares, S., Moreira, L., Matias, S., Itskov, P.M., Correia, P.A., et al. (2015). Bonsai: an event-based framework for processing and controlling data streams. Front Neuroinform 9, 7. 10.3389/fninf.2015.00007.

43. Lu, J., Li, C., Singh-Alvarado, J., Zhou, Z.C., Fröhlich, F., Mooney, R., and Wang, F. (2018). MIN1PIPE: A Miniscope 1-Photon-Based Calcium Imaging Signal Extraction Pipeline. CellReports 23, 3673–3684. 10.1016/j.celrep.2018.05.062.

44. Sheintuch, L., Rubin, A., Brande-Eilat, N., Geva, N., Sadeh, N., Pinchasof, O., and Ziv, Y. (2017). Tracking the Same Neurons across Multiple Days in Ca2+ Imaging Data. Cell Reports 21, 1102–1115. 10.1016/j.celrep.2017.10.013.

45. Bankhead, P., Loughrey, M.B., Fernández, J.A., Dombrowski, Y., McArt, D.G., Dunne, P.D., McQuaid, S., Gray, R.T., Murray, L.J., Coleman, H.G., et al. (2017). QuPath: Open source software for digital pathology image analysis. Sci. Rep. 7, 16878. 10.1038/s41598-017-17204-5.

