## Supporting information for "Threat-dependent scaling of prelimbic dynamics to enhance fear representation"

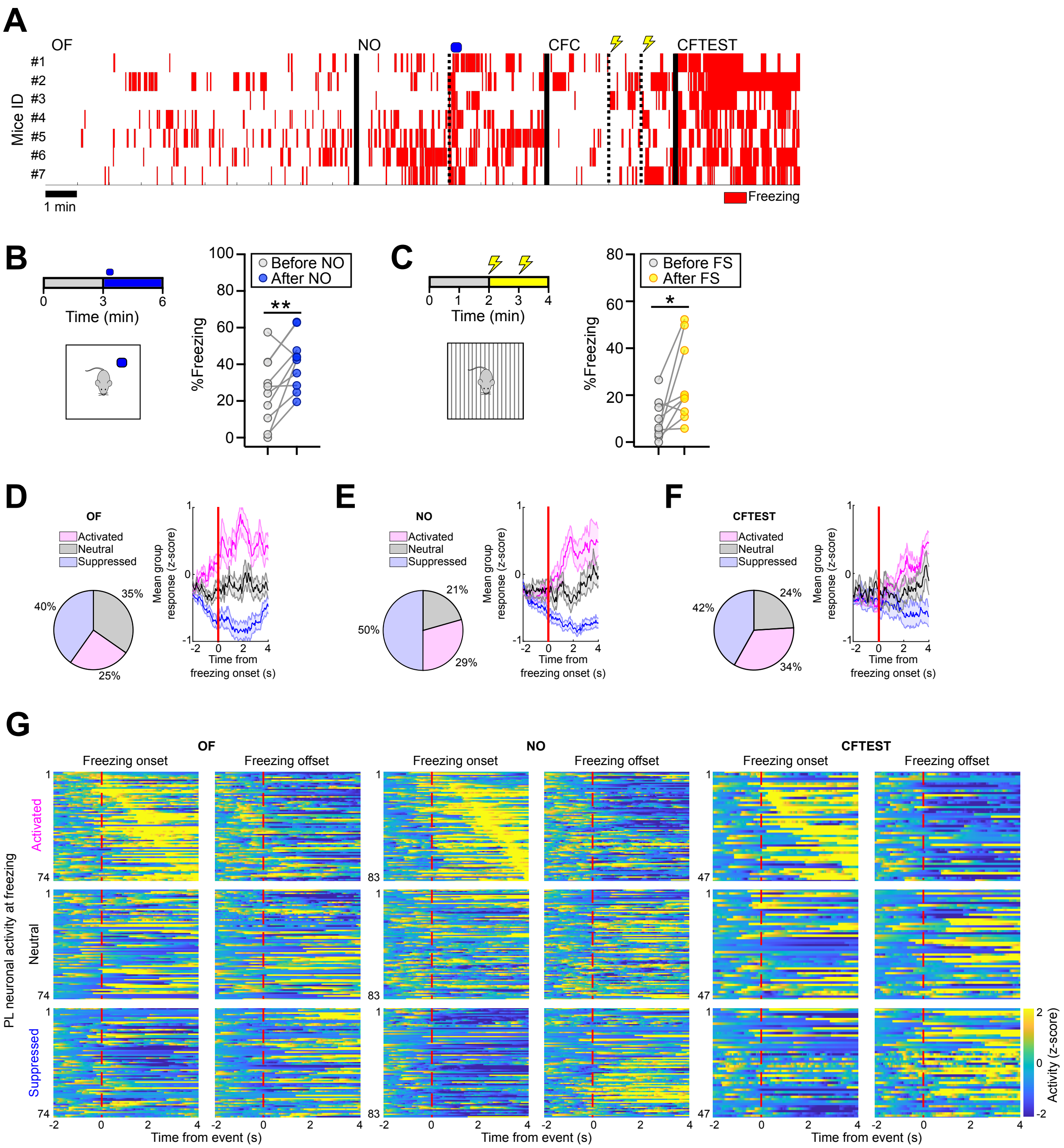

**Figure S1. Behavioral analysis and PL single-cell encoding of freezing**. **(A)** Time course of freezing episodes during Open Field (OF), Novel Object (NO), Contextual Fear Conditioning (CFC), and Contextual Fear Test (CFTEST). **(B)** Freezing scoring during the Novel Object (NO) test, before and after the introduction of the object. **(C)** Freezing scoring during Contextual Fear Conditioning (CFC) training before and after the foot shocks (FS). **(D-F)** Pie charts showing the percentage of all imaged PL cells classified as activated, suppressed, or neutral to freezing during Open Field (OF, **(D)**, N=1052 cells), Novel Object (NO, **(E)**, N=1078 cells), and Contextual Fear Conditioning TEST (CFTEST, **(F)**, N=695 cells). Mean group response to freezing onset (red line) is also shown (data represent mean ± SEM). **(G)** Heatmaps showing normalized activity of cells aligned to freezing (dotted red line) onset and offset.

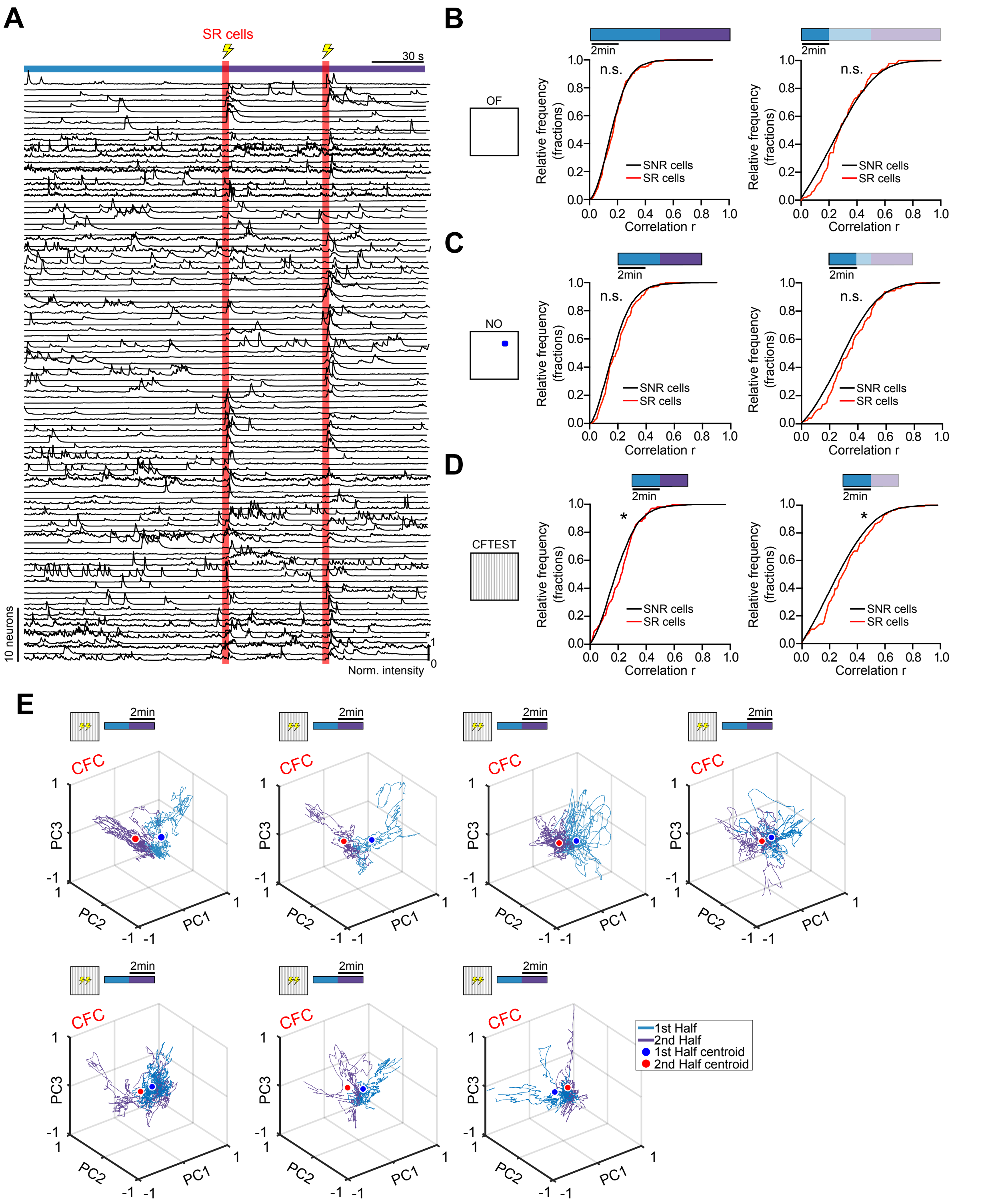

**Figure S2. Higher correlated activity of Shock-responding cells (SR) cells vs Shock-nonresponding cells (SNR) cells following Contextual Fear Conditioning (CFC). (A)** Calcium traces of 84 SR cells recorded during CFC (total N=104 cells, from 9 mice). **(B)** Relative frequency plots of correlation of activity during Open Field (OF), **(C)** Novel Object (NO), and **(D)** Contextual Fear Conditioning Test (CFTEST). Left panels consider the whole task period while right panels show only the first 2 min of each task. **(E)** Representative PCA clustering of Prelimbic cortex (PL) neuronal activity across time during CFC; red and blue dots represent the cluster centroids. *P<0.05; n.s.= P>0.05.

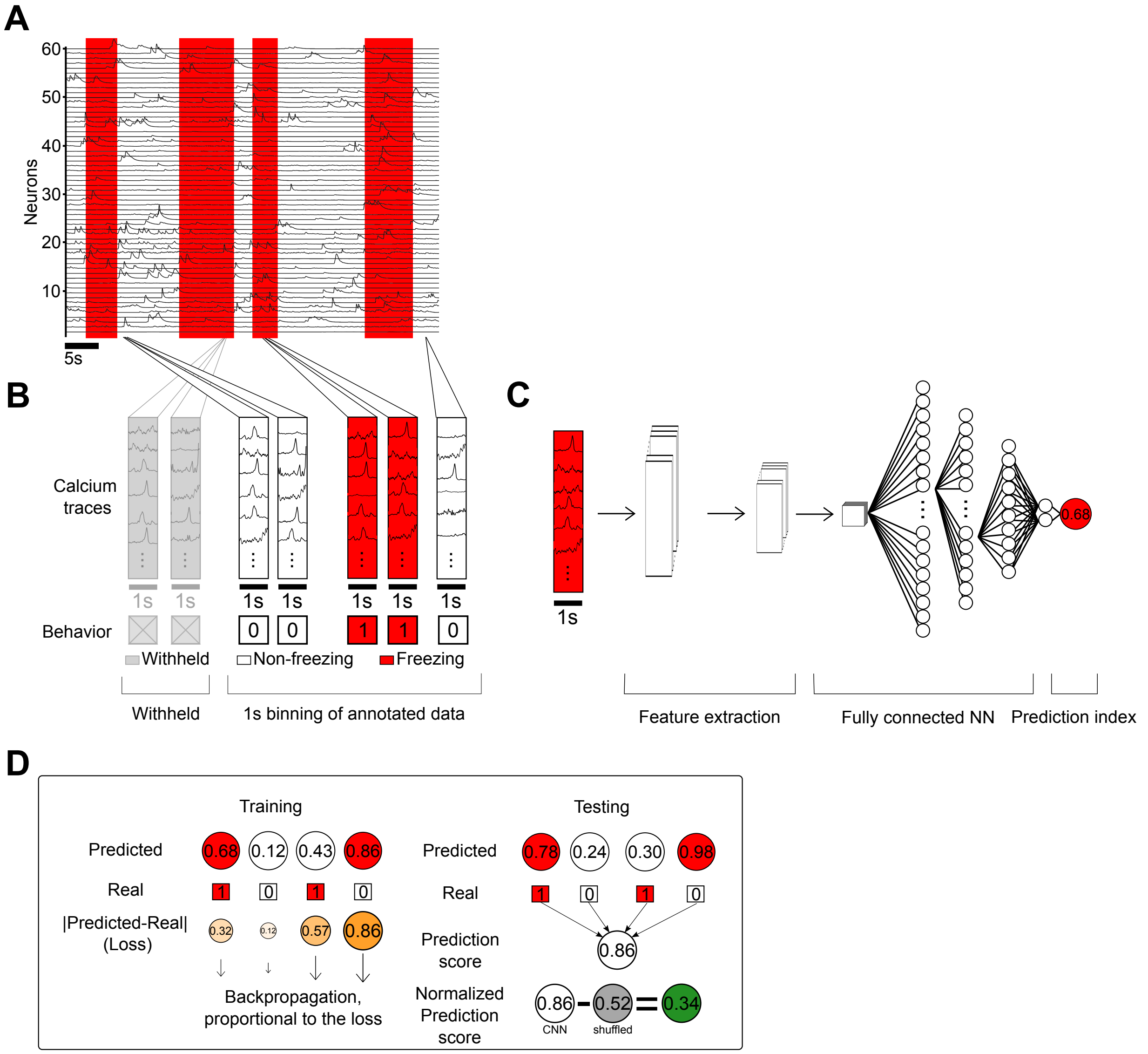

**Figure S3. Convolutional neural network (CNN) design. (A)** Calcium traces of the 60 most active neurons are annotated according to animal behavior: freezing (red) or no freezing (white). **(B)** These traces are then split into one second bouts of annotated data that are used as CNN input (at 20 fps, the input is a 60-by-20 matrix). Some of the data is withheld during training to be used during the test. **(C)** Each bout of annotated data is inputted into the CNN: it first goes through feature extraction steps, then fully connected layers (see methods). The CNN outputs a prediction of the animal's behavior (where 0 is non-freezing and 1 is freezing) in the form of a 'prediction index'. The prediction ranges from 0 to 1. Anything above 0.5 is considered a 'freezing' guess, anything below 0.5 is considered a 'non-freezing' guess. **(D)** During training (left), every guess is compared to the ground truth values, and the CNN updates its weights via backpropagation to improve its predictions. During testing (right), the CNN only evaluates bouts that were withheld during training. The predictions are compared to the real values. They are regrouped by animal and experiment, and weighted to give equal importance to no-freezing and freezing data to obtain prediction scores. The train/test split is repeated 100 times such that every activity bout occurs at least 20 times in a test set. This is repeated with randomly shuffled annotations to estimate chance performance and obtain a normalized prediction score.

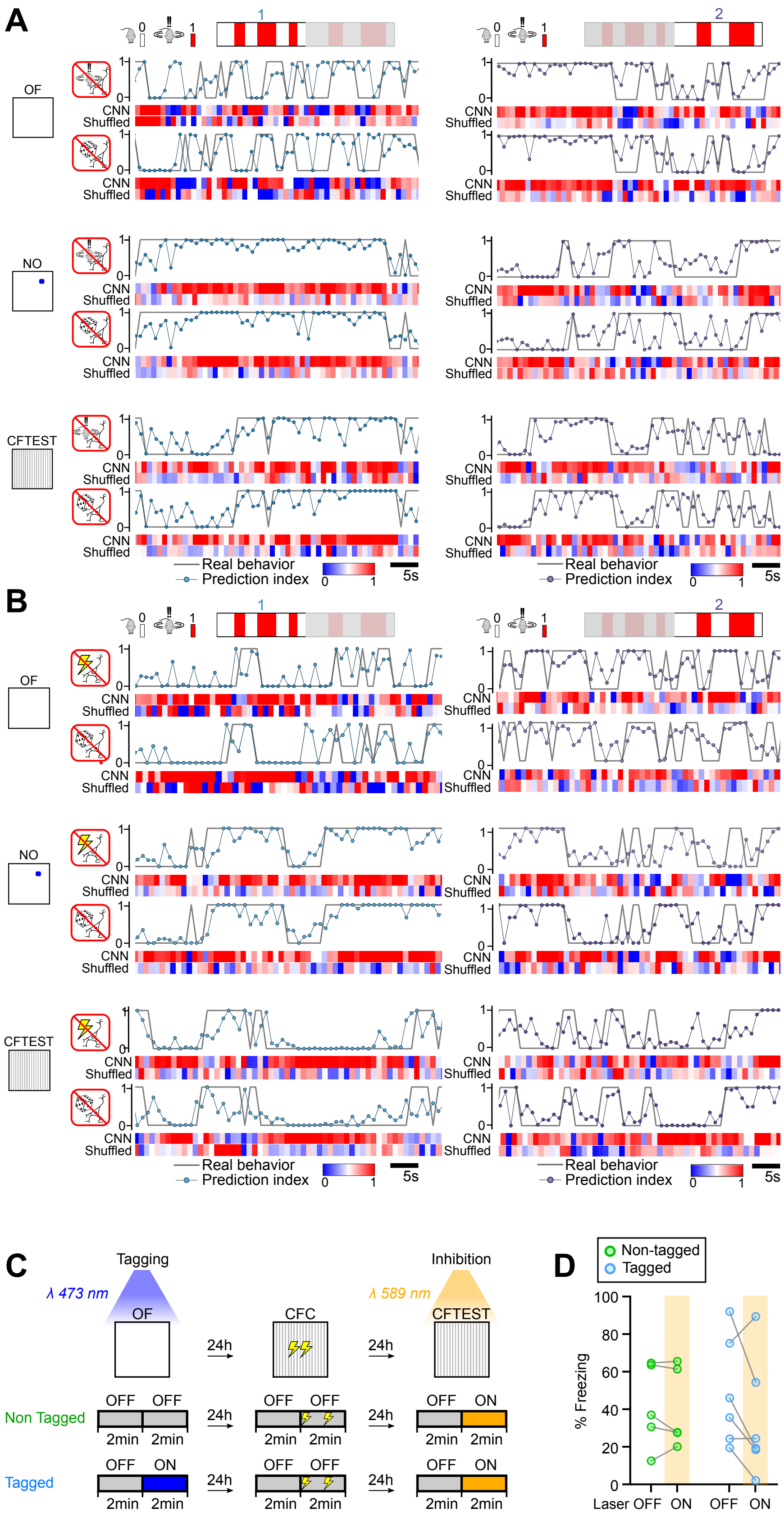
A
B
C
D
1
2
OF
NO
CFTEST
CNN
Shuffled
Real behavior
Prediction index
5s
Tagging
Inhibition
λ 473 nm
λ 589 nm
CFC
24h
OFF
ON
2min
Non Tagged
Tagged
Non-tagged
% Freezing
Laser OFF ON OFF ON

**Figure S4. (A-B)** Examples of prediction scores for one mouse in different tasks and periods excluding either freezing activated or random cells **(A)** and excluding either shock-responding or random cells **(B)**. Top: Prediction indices obtained for every 1s of input data, compared to the real behavior. Different values are assigned to the 2 behaviors, either real in full lines or predicted in dotted lines: 1 is freezing, and 0 is non-freezing. Bottom, performance of CNN with either real or shuffled behavioral annotations. Red blue scale: 1 perfect guess, 0 completely opposite guess. **(C)** Schematics of experimental setup for control FLiCRE experiments. Following viral injection and optic fiber implantation, mice underwent OF while blue light was continuously delivered to the PL during the last 2 minutes of the task, enabling the expression of inhibitory opsin eNpHR3.0 in active neurons only for mice belonging to the “tagged” group. The following day, mice were trained in CFC. One day later during CFTEST, all mice PL were illuminated using continuous yellow light during the last 2 min to inhibit activity of neurons expressing eNpHR3.0. **(D)** Percentage of time spent in freezing during CFTEST. Yellow shaded area represents the period when the yellow laser was on.

**Supplementary Table 1.**

| MAIN FIGURES | | | | | | |
|---|---|---|---|---|---|---|
| FIG | TEST | VARIABLE | GROUPS | N | STATISTIC | P |
| 1C | Mixed effects model (REML) | %Freezing | OF | 10 | F (1.381,14.73) = 35,07 | <0.0001 |
| | | | NO | 10 | | |
| | | | CFC | 9 | | |
| | | | CFTest | 7 | | |
| | Tukey's multiple comparison test | %Freezing | OF vs NO | | | 0.0021 |
| | | %Freezing | OF vs CFC | | | 0.5523 |
| | | %Freezing | OF vs CFTEST | | | <0.0001 |
| | | %Freezing | NO vs CFC | | | 0.1505 |
| | | %Freezing | NO vs CFTEST | | | 0.0381 |
| | | %Freezing | CFC vs CFTEST | | | <0.0001 |
| 1M | Kolmogorov-Smirnov | Frequency distribution of correlation r value | CFC last 2 min; Shock-responding cells vs Shock-nonresponding cells | | D=0.1554 | <0.0001 |
| 1N | Kolmogorov-Smirnov | Frequency distribution of correlation r value | CFTEST last 2 min; Shock-responding cells vs Shock-nonresponding cells | | D=0.1898 | 0.001 |
| 1O | Kolmogorov-Smirnov | Frequency distribution of correlation r value | CFC first 2 min; Shock-responding cells vs Shock-nonresponding cells | | D=0.1064 | <0.0001 |
| 1P | Kolmogorov-Smirnov | Frequency distribution of correlation r value | OF last 2 min; Shock-responding cells vs Shock-nonresponding cells | | D=0.1299 | 0.0764 |
| 1Q | Kolmogorov-Smirnov | Frequency distribution of correlation r value | NO last 2 min; Shock-responding cells vs Shock-nonresponding cells | | D=0.1027 | 0.1446 |
| 2C | Mixed effects model (REML) | Centroid Euclidean distance between PCA-obtained clusters | OF | 10 | F(1.846,13.54)=14.74 | 0.0005 |
| | | | NO | 10 | | |
| | | | CFC | 9 | | |
| | | | CFTEST | 7 | | |
| | Tukey's multiple comparison test | Centroid Euclidean distance between PCA-obtained clusters | OF vs NO | | | 0.8428 |
| | | Centroid Euclidean distance between PCA-obtained clusters | OF vs CFC | | | 0.0079 |
| | | Centroid Euclidean distance between PCA-obtained clusters | OF vs CFTEST | | | 0.256 |
| | | Centroid Euclidean distance between PCA-obtained clusters | NO vs CFC | | | 0.0028 |
| | | Centroid Euclidean distance between PCA-obtained clusters | NO vs CFTEST | | | 0.3225 |
| | | Centroid Euclidean distance between PCA-obtained clusters | CFC vs CFTEST | | | 0.0187 |
| 2H | Kruskal-Wallis test | Centroid Euclidean distance between PCA-obtained clusters | PL | 10 | H(2)=11.72 | 0.0006 |
| | | | LDn | 8 | | |
| | | | RSC | 5 | | |
| | Dunn's multiple comparison test | Centroid Euclidean distance between PCA-obtained clusters | PL vs LDn | | | 0.0112 |
| | | | PL vs RSC | | | 0.011 |
| | | | LDn vs RSC | | | >0.9999 |
| 2J | Wilcoxon signed rank test | Prediction score (CNN - Shuffled) | OF1 (CNN - Shuffled) | 10 | W(10) = 55 | 0.002 |
| | Wilcoxon signed rank test | Prediction score (CNN - Shuffled) | OF2 (CNN - Shuffled) | 10 | W(10) = 55 | 0.002 |
| | Wilcoxon signed rank test | Prediction score (CNN - Shuffled) | NO1 (CNN - Shuffled) | 10 | W(10) = 55 | 0.002 |
| | Wilcoxon signed rank test | Prediction score (CNN - Shuffled) | NO2 (CNN - Shuffled) | 10 | W(10) = 53 | 0.0039 |
| | Wilcoxon signed rank test | Prediction score (CNN - Shuffled) | CFC1 (CNN - Shuffled) | 9 | W(9) = 45 | 0.0039 |
| | Wilcoxon signed rank test | Prediction score (CNN - Shuffled) | CFC2 (CNN - Shuffled) | 9 | W(9) = 45 | 0.0039 |

| | | | | | | |
|---|---|---|---|---|---|---|
| | Wilcoxon signed rank test | Prediction score (CNN - Shuffled) | CFTEST1 (CNN - Shuffled) | 7 | W(7) = 28 | 0.0156 |
| | Wilcoxon signed rank test | Prediction score (CNN - Shuffled) | CFTEST2 (CNN - Shuffled) | 7 | W(7) = 28 | 0.0156 |
| | Wilcoxon matched-pairs signed rank test | Normalized prediction score | OF1 vs OF2 | 10 | W(10) = -13 | 0.5566 |
| | Wilcoxon matched-pairs signed rank test | Normalized prediction score | NO1 vs NO2 | 10 | W(10) = -27 | 0.1934 |
| | Wilcoxon matched-pairs signed rank test | Normalized prediction score | CFC1 vs CFC2 | 9 | W(9) = 43 | 0.0078 |
| | Wilcoxon matched-pairs signed rank test | Normalized prediction score | CFTEST1 vs CFTEST2 | 7 | W(7) = -4 | 0.8125 |
| 2L | Kruskal-Wallis test | Normalized prediction scores | PL | 10 | H(2)=15.15 | <0.0001 |
| | | | LDn | 8 | | |
| | | | RSC | 4 | | |
| | Dunn's multiple comparison test | Normalized prediction scores | PL vs LDn | | | 0.0016 |
| | | | PL vs RSC | | | 0.011 |
| | | | LDn vs RSC | | | >0.9999 |
| | Wilcoxon signed rank test | Prediction score (CNN - Shuffled) | OF PL | 10 | W(10) = 55 | 0.002 |
| | Wilcoxon signed rank test | Prediction score (CNN - Shuffled) | OF LDn | 8 | W(8) = 34 | 0.0156 |
| | Wilcoxon signed rank test | Prediction score (CNN - Shuffled) | OF RSC | 4 | W(4) = 10 | 0.125 |
| 3A (top left) | Wilcoxon signed rank test | Prediction score (CNN - Shuffled) | OF1, regular training | 10 | W(10) = 55 | 0.002 |
| | Wilcoxon signed rank test | Prediction score (CNN - Shuffled) | OF2, regular training | 10 | W(10) = 55 | 0.002 |
| | Wilcoxon signed rank test | Prediction score (CNN - Shuffled) | OF1, Freezing cells excluded | 10 | W(10) = 55 | 0.002 |
| | Wilcoxon signed rank test | Prediction score (CNN - Shuffled) | OF2, Freezing cells excluded | 10 | W(10) = 55 | 0.002 |
| | Wilcoxon signed rank test | Prediction score (CNN - Shuffled) | OF1, Random cells excluded | 10 | W(10) = 55 | 0.002 |
| | Wilcoxon signed rank test | Prediction score (CNN - Shuffled) | OF2, Random cells excluded | 10 | W(10) = 55 | 0.002 |
| | Wilcoxon matched-pairs signed rank test | Normalized prediction scores | OF1 vs OF2, regular training | 10 | W(10) = -27 | 0.1934 |
| | Wilcoxon matched-pairs signed rank test | Normalized prediction scores | OF1 vs OF2, Freezing cells excluded | 10 | W(10) = 7 | 0.7695 |
| | Wilcoxon matched-pairs signed rank test | Normalized prediction scores | OF1 vs OF2,, Random cells excluded | 10 | W(10) = 5 | 0.8457 |
| 3A (top right) | Wilcoxon signed rank test | Prediction score (CNN - Shuffled) | NO1, regular training | 10 | W(10) = 55 | 0.002 |
| | Wilcoxon signed rank test | Prediction score (CNN - Shuffled) | NO2, regular training | 10 | W(10) = 53 | 0.0039 |
| | Wilcoxon signed rank test | Prediction score (CNN - Shuffled) | NO1, Freezing cells excluded | 10 | W(10) = 55 | 0.002 |
| | Wilcoxon signed rank test | Prediction score (CNN - Shuffled) | NO2, Freezing cells excluded | 10 | W(10) = 55 | 0.002 |
| | Wilcoxon signed rank test | Prediction score (CNN - Shuffled) | NO1, Random cells excluded | 10 | W(10) = 55 | 0.002 |
| | Wilcoxon signed rank test | Prediction score (CNN - Shuffled) | NO2, Random cells excluded | 10 | W(10) = 55 | 0.002 |
| | Wilcoxon matched-pairs signed rank test | Normalized prediction scores | NO1 vs NO2, regular training | 10 | W(10) = -13 | 0.5566 |
| | Wilcoxon matched-pairs signed rank test | Normalized prediction scores | NO1 vs NO2, Freezing cells excluded | 10 | W(10) = -29 | 0.1602 |
| | Wilcoxon matched-pairs signed rank test | Normalized prediction scores | NO1 vs NO2, Random cells excluded | 10 | W(10) = -17 | 0.4316 |
| 3A (bottom left) | Wilcoxon signed rank test | Prediction score (CNN - Shuffled) | CFC1, regular training | 9 | W(9) = 45 | 0.0039 |
| | Wilcoxon signed rank test | Prediction score (CNN - Shuffled) | CFC2, regular training | 9 | W(9) = 45 | 0.0039 |
| | Wilcoxon signed rank test | Prediction score (CNN - Shuffled) | CFC1, Freezing cells excluded | 9 | W(9) = 41 | 0.0117 |
| | Wilcoxon signed rank test | Prediction score (CNN - Shuffled) | CFC2, Freezing cells excluded | 9 | W(9) = 45 | 0.0039 |
| | Wilcoxon signed rank test | Prediction score (CNN - Shuffled) | CFC1, Random cells excluded | 9 | W(9) = 45 | 0.0039 |
| | Wilcoxon signed rank test | Prediction score (CNN - Shuffled) | CFC2, Random cells excluded | 9 | W(9) = 45 | 0.0039 |

| | | | | | | |
|---|---|---|---|---|---|---|
| | Wilcoxon matched-pairs signed rank test | Normalized prediction scores | CFC1 vs CFC2, regular training | 9 | W(9) = 43 | 0.0078 |
| | Wilcoxon matched-pairs signed rank test | Normalized prediction scores | CFC1 vs CFC2, Freezing cells excluded | 9 | W(9) = 39 | 0.0195 |
| | Wilcoxon matched-pairs signed rank test | Normalized prediction scores | CFC1 vs CFC2, Random cells excluded | 9 | W(9) = 43 | 0.0078 |
| 3A (bottom right) | Wilcoxon signed rank test | Prediction score (CNN - Shuffled) | CFTEST1, regular training | 7 | W(7) = 28 | 0.0156 |
| | Wilcoxon signed rank test | Prediction score (CNN - Shuffled) | CFTEST2, regular training | 7 | W(7) = 28 | 0.0156 |
| | Wilcoxon signed rank test | Prediction score (CNN - Shuffled) | CFTEST1, Freezing cells excluded | 7 | W(7) = 28 | 0.0156 |
| | Wilcoxon signed rank test | Prediction score (CNN - Shuffled) | CFTEST2, Freezing cells excluded | 7 | W(7) = 28 | 0.0156 |
| | Wilcoxon signed rank test | Prediction score (CNN - Shuffled) | CFTEST1, Random cells excluded | 7 | W(7) = 28 | 0.0156 |
| | Wilcoxon signed rank test | Prediction score (CNN - Shuffled) | CFTEST2, Random cells excluded | 7 | W(7) = 28 | 0.0156 |
| | Wilcoxon matched-pairs signed rank test | Normalized prediction scores | CFTEST1 vs CFTEST2, regular training | 7 | W(7) = -20 | 0.1094 |
| | Wilcoxon matched-pairs signed rank test | Normalized prediction scores | CFTEST1 vs CFTEST2, Freezing cells excluded | 7 | W(7) = 4 | 0.8125 |
| | Wilcoxon matched-pairs signed rank test | Normalized prediction scores | CFTEST1 vs CFTEST2, Random cells excluded | 7 | W(7) = -22 | 0.0781 |
| 3C left | Wilcoxon signed rank test | Prediction score (CNN - Shuffled) | CFC1, regular training | 8 | W(8) = 30 | 0.0391 |
| | Wilcoxon signed rank test | Prediction score (CNN - Shuffled) | CFC2, regular training | 8 | W(8) = 34 | 0.0156 |
| | Wilcoxon signed rank test | Prediction score (CNN - Shuffled) | CFC1, Freezing cells excluded | 8 | W(8) = 24 | 0.1094 |
| | Wilcoxon signed rank test | Prediction score (CNN - Shuffled) | CFC2, Freezing cells excluded | 8 | W(8) = 34 | 0.0156 |
| | Wilcoxon signed rank test | Prediction score (CNN - Shuffled) | CFC1, Random cells excluded | 8 | W(8) = 24 | 0.1094 |
| | Wilcoxon signed rank test | Prediction score (CNN - Shuffled) | CFC2, Random cells excluded | 8 | W(8) = 30 | 0.0391 |
| | Wilcoxon matched-pairs signed rank test | Normalized prediction scores | CFC1 vs CFC2, regular training | 8 | W(8)=18 | 0.25 |
| | Wilcoxon matched-pairs signed rank test | Normalized prediction scores | CFC1 vs CFC2, Freezing cells excluded | 8 | W(8)=28 | 0.0547 |
| | Wilcoxon matched-pairs signed rank test | Normalized prediction scores | CFC1 vs CFC2, Random cells excluded | 8 | W(8)=-2 | 0.9453 |
| 3C right | Wilcoxon signed rank test | Prediction score (CNN - Shuffled) | CFC1, regular training | 5 | W(5) = 13 | 0.125 |
| | Wilcoxon signed rank test | Prediction score (CNN - Shuffled) | CFC2, regular training | 5 | W(5) = 13 | 0.125 |
| | Wilcoxon signed rank test | Prediction score (CNN - Shuffled) | CFC1, Freezing cells excluded | 5 | W(5) = 13 | 0.125 |
| | Wilcoxon signed rank test | Prediction score (CNN - Shuffled) | CFC2, Freezing cells excluded | 5 | W(5) = 11 | 0.1875 |
| | Wilcoxon signed rank test | Prediction score (CNN - Shuffled) | CFC1, Random cells excluded | 5 | W(5) = 7 | 0.4375 |
| | Wilcoxon signed rank test | Prediction score (CNN - Shuffled) | CFC2, Random cells excluded | 5 | W(5) = 13 | 0.125 |
| | Wilcoxon matched-pairs signed rank test | Normalized prediction scores | CFC1 vs CFC2, regular training | 5 | W(5) = 1 | >0.9999 |
| | Wilcoxon matched-pairs signed rank test | Normalized prediction scores | CFC1 vs CFC2, Freezing cells excluded | 5 | W(5) = 7 | 0.4375 |
| | Wilcoxon matched-pairs signed rank test | Normalized prediction scores | CFC1 vs CFC2, Random cells excluded | 5 | W(5) = 9 | 0.3125 |
| 3D (top left) | Wilcoxon signed rank test | Prediction score (CNN - Shuffled) | OF1, regular training | 9 | W(9) = 45 | 0.0039 |
| | Wilcoxon signed rank test | Prediction score (CNN - Shuffled) | OF2, regular training | 9 | W(9) = 45 | 0.0039 |
| | Wilcoxon signed rank test | Prediction score (CNN - Shuffled) | OF1, Shock cells excluded | 9 | W(9) = 43 | 0.0078 |
| | Wilcoxon signed rank test | Prediction score (CNN - Shuffled) | OF2, Shock cells excluded | 9 | W(9) = 41 | 0.0117 |
| | Wilcoxon signed rank test | Prediction score (CNN - Shuffled) | OF1, Random cells excluded | 9 | W(9) = 41 | 0.0117 |
| | Wilcoxon signed rank test | Prediction score (CNN - Shuffled) | OF2, Random cells excluded | 9 | W(9) = 37 | 0.0273 |
| | Wilcoxon matched-pairs signed rank test | Normalized prediction scores | OF1 vs OF2, regular training | 9 | W(10) = -19 | 0.3008 |

| | | | | | | |
|---|---|---|---|---|---|---|
| | Wilcoxon matched-pairs signed rank test | Normalized prediction scores | OF1 vs OF2, Shock cells excluded | 9 | W(9) = -13 | 0.4951 |
| | Wilcoxon matched-pairs signed rank test | Normalized prediction scores | OF1 vs OF2,, Random cells excluded | 9 | W(9) = -25 | 0.1641 |
| 3D (top right) | Wilcoxon signed rank test | Prediction score (CNN - Shuffled) | NO1, regular training | 9 | W(10) = 45 | 0.0039 |
| | Wilcoxon signed rank test | Prediction score (CNN - Shuffled) | NO2, regular training | 9 | W(10) = 43 | 0.0078 |
| | Wilcoxon signed rank test | Prediction score (CNN - Shuffled) | NO1, Shock cells excluded | 9 | W(9) = 39 | 0.0195 |
| | Wilcoxon signed rank test | Prediction score (CNN - Shuffled) | NO2, Shock cells excluded | 9 | W(9) = 39 | 0.0195 |
| | Wilcoxon signed rank test | Prediction score (CNN - Shuffled) | NO1, Random cells excluded | 9 | W(9) = 39 | 0.0195 |
| | Wilcoxon signed rank test | Prediction score (CNN - Shuffled) | NO2, Random cells excluded | 9 | W(9) = 45 | 0.0039 |
| | Wilcoxon matched-pairs signed rank test | Normalized prediction scores | NO1 vs NO2, regular training | 9 | W(9) = -21 | 0.25 |
| | Wilcoxon matched-pairs signed rank test | Normalized prediction scores | NO1 vs NO2, Shock cells excluded | 9 | W(9) = 3 | 0.9102 |
| | Wilcoxon matched-pairs signed rank test | Normalized prediction scores | NO1 vs NO2, Random cells excluded | 9 | W(9) = -3 | 0.9102 |
| 3D (bottom left) | Wilcoxon signed rank test | Prediction score (CNN - Shuffled) | CFC1, regular training | 9 | W(9) = 45 | 0.0039 |
| | Wilcoxon signed rank test | Prediction score (CNN - Shuffled) | CFC2, regular training | 9 | W(9) = 45 | 0.0039 |
| | Wilcoxon signed rank test | Prediction score (CNN - Shuffled) | CFC1, Shock cells excluded | 9 | W(9) = 45 | 0.0039 |
| | Wilcoxon signed rank test | Prediction score (CNN - Shuffled) | CFC2, Shock cells excluded | 9 | W(9) = 45 | 0.0039 |
| | Wilcoxon signed rank test | Prediction score (CNN - Shuffled) | CFC1, Random cells excluded | 9 | W(9) = 37 | 0.0273 |
| | Wilcoxon signed rank test | Prediction score (CNN - Shuffled) | CFC2, Random cells excluded | 9 | W(9) = 45 | 0.0039 |
| | Wilcoxon matched-pairs signed rank test | Normalized prediction scores | CFC1 vs CFC2, regular training | 9 | W(9) = 43 | 0.0078 |
| | Wilcoxon matched-pairs signed rank test | Normalized prediction scores | CFC1 vs CFC2, Shock cells excluded | 9 | W(9) = 35 | 0.0391 |
| | Wilcoxon matched-pairs signed rank test | Normalized prediction scores | CFC1 vs CFC2, Random cells excluded | 9 | W(9) = 39 | 0.0195 |
| 3D (bottom right) | Wilcoxon signed rank test | Prediction score (CNN - Shuffled) | CFTEST1, regular training | 7 | W(7) = 28 | 0.0156 |
| | Wilcoxon signed rank test | Prediction score (CNN - Shuffled) | CFTEST2, regular training | 7 | W(7) = 28 | 0.0156 |
| | Wilcoxon signed rank test | Prediction score (CNN - Shuffled) | CFTEST1, Shock cells excluded | 7 | W(7) = 28 | 0.0156 |
| | Wilcoxon signed rank test | Prediction score (CNN - Shuffled) | CFTEST2, Shock cells excluded | 7 | W(7) = 28 | 0.0156 |
| | Wilcoxon signed rank test | Prediction score (CNN - Shuffled) | CFTEST1, Random cells excluded | 7 | W(7) = 26 | 0.0312 |
| | Wilcoxon signed rank test | Prediction score (CNN - Shuffled) | CFTEST2, Random cells excluded | 7 | W(7) = 26 | 0.0312 |
| | Wilcoxon matched-pairs signed rank test | Normalized prediction scores | CFTEST1 vs CFTEST2, regular training | 7 | W(7) = -20 | 0.1094 |
| | Wilcoxon matched-pairs signed rank test | Normalized prediction scores | CFTEST1 vs CFTEST2, Shock cells excluded | 7 | W(7) = -12 | 0.375 |
| | Wilcoxon matched-pairs signed rank test | Normalized prediction scores | CFTEST1 vs CFTEST2, Random cells excluded | 7 | W(7) = 4 | 0.8125 |
| 3F left | Wilcoxon signed rank test | Prediction score (CNN - Shuffled) | CFC1, regular training | 8 | W(8) = 30 | 0.0391 |
| | Wilcoxon signed rank test | Prediction score (CNN - Shuffled) | CFC2, regular training | 8 | W(8) = 34 | 0.0156 |
| | Wilcoxon signed rank test | Prediction score (CNN - Shuffled) | CFC1, Freezing cells excluded | 8 | W(8) = 10 | 0.5469 |
| | Wilcoxon signed rank test | Prediction score (CNN - Shuffled) | CFC2, Freezing cells excluded | 8 | W(8) = 24 | 0.1094 |
| | Wilcoxon signed rank test | Prediction score (CNN - Shuffled) | CFC1, Random cells excluded | 8 | W(8) = 26 | 0.0781 |
| | Wilcoxon signed rank test | Prediction score (CNN - Shuffled) | CFC2, Random cells excluded | 8 | W(8) = 26 | 0.0781 |
| | Wilcoxon matched-pairs signed rank test | Normalized prediction scores | CFC1 vs CFC2, regular training | 8 | W(8)=14 | 0.25 |

| | | | | | | |
|---|---|---|---|---|---|---|
| | Wilcoxon matched-pairs signed rank test | Normalized prediction scores | CFC1 vs CFC2, Freezing cells excluded | 8 | W(8)=28 | 0.3828 |
| | Wilcoxon matched-pairs signed rank test | Normalized prediction scores | CFC1 vs CFC2, Random cells excluded | 8 | W(8)=-2 | 0.9453 |
| 3F right | Wilcoxon signed rank test | Prediction score (CNN - Shuffled) | CFC1, regular training | 5 | W(5) = 13 | 0.125 |
| | Wilcoxon signed rank test | Prediction score (CNN - Shuffled) | CFC2, regular training | 5 | W(5) = 13 | 0.125 |
| | Wilcoxon signed rank test | Prediction score (CNN - Shuffled) | CFC1, Freezing cells excluded | 5 | W(5) = 13 | 0.125 |
| | Wilcoxon signed rank test | Prediction score (CNN - Shuffled) | CFC2, Freezing cells excluded | 5 | W(5) = 11 | 0.1875 |
| | Wilcoxon signed rank test | Prediction score (CNN - Shuffled) | CFC1, Random cells excluded | 5 | W(5) = 13 | 0.125 |
| | Wilcoxon signed rank test | Prediction score (CNN - Shuffled) | CFC2, Random cells excluded | 5 | W(5) = 11 | 0.1875 |
| | Wilcoxon matched-pairs signed rank test | Normalized prediction scores | CFC1 vs CFC2, regular training | 5 | W(5) = 1 | >0.9999 |
| | Wilcoxon matched-pairs signed rank test | Normalized prediction scores | CFC1 vs CFC2, Freezing cells excluded | 5 | W(5) = -7 | 0.4375 |
| | Wilcoxon matched-pairs signed rank test | Normalized prediction scores | CFC1 vs CFC2, Random cells excluded | 5 | W(5) = -13 | 0.125 |
| 4E left | Mann Whitney test | CaM-uTEVp+ neurons / mm² | Tagged before foot shocks vs Tagged after foot shocks | | U=28 | >0.9999 |
| | | | Tagged before foot shocks | 7 | | |
| | | | Tagged after foot shocks | 8 | | |
| 4E middle | Mann Whitney test | mCherry+ neurons / mm² | Tagged before foot shocks vs Tagged after foot shocks | | U=22 | 0.5358 |
| | | | Tagged before foot shocks | 7 | | |
| | | | Tagged after foot shocks | 8 | | |
| 4E right | Mann Whitney test | mCherry+ / CaM-uTEVp+ neurons (%) | Tagged before foot shocks vs Tagged after foot shocks | | U=14 | 0.1206 |
| | | | Tagged before foot shocks | 7 | | |
| | | | Tagged after foot shocks | 8 | | |
| 4F | Wilcoxon matched-pairs signed rank test | %Freezing | Laser OFF vs Laser ON, Tagged before foot shocks | 7 | W(7)=-8 | 0.5781 |
| | Wilcoxon matched-pairs signed rank test | %Freezing | Laser OFF vs Laser ON, Tagged after foot shocks | 8 | W(8)=-36 | 0.0078 |
| **SUPPLEMENTARY FIGURES** | | | | | | |
| **FIG** | **TEST** | **VARIABLE** | **GROUPS** | **N** | **STATISTIC** | **P** |
| S1B | Wilcoxon matched-pairs signed rank test | %Freezing | NO before object | 10 | W(10)=49 | 0.0098 |
| | | | NO after object | | | |
| S1C | Wilcoxon matched-pairs signed rank test | %Freezing | CFC before shocks | 9 | W(9)=41 | 0.0117 |
| | | | CFC after shocks | | | |
| S2B left | Kolmogorov-Smirnov | Frequency distribution of correlation r value | OF; Shock-responding cells vs Shock-nonresponding cells | | D=0.06732 | 0.5738 |
| S2B right | Kolmogorov-Smirnov | Frequency distribution of correlation r value | OF first 2 min; Shock-responding cells vs Shock-nonresponding cells | | D=0.1305 | 0.0741 |
| S2C left | Kolmogorov-Smirnov | Frequency distribution of correlation r value | NO; Shock-responding cells vs Shock-nonresponding cells | | D=0.1033 | 0.1032 |
| S2C right | Kolmogorov-Smirnov | Frequency distribution of correlation r value | NO first 2min; Shock-responding cells vs Shock-nonresponding cells | | D=0.1201 | 0.055 |
| S2D left | Kolmogorov-Smirnov | Frequency distribution of correlation r value | CFTEST; Shock-responding cells vs Shock-nonresponding cells | | D=0.1484 | 0.019 |
| S2D right | Kolmogorov-Smirnov | Frequency distribution of correlation r value | CFTEST first 2 min; Shock-responding cells vs Shock-nonresponding cells | | D=0.1407 | 0.0294 |
| S4D | Wilcoxon matched-pairs signed rank test | %Freezing | Laser OFF vs Laser ON, Non-tagged | 5 | W(5)=-5 | 0.625 |
| | Wilcoxon matched-pairs signed rank test | %Freezing | Laser OFF vs Laser ON, Tagged OF last 2 min | 6 | W(6)=-15 | 0.1562 |
